# Mammalian MemPrep establishes the lipid composition of ER membranes in HEK293T cells

**DOI:** 10.64898/2026.03.20.713196

**Authors:** Aamna Jain, Alexander von der Malsburg, Cynthia Alsayyah, Claudia Götz, Mohamed Elmofty, John Reinhard, Per Haberkant, Volkhard Helms, Joseph H. Lorent, Robert Ernst

## Abstract

The endoplasmic reticulum (ER) forms a dynamic network of sheets and tubules, whose molecular lipid composition remains incompletely defined. Using an optimized MemPrep workflow, we establish a high-confidence lipidome of the mammalian ER and selectively enrich membrane vesicles originating from ER tubules as a major ER subdomain. Quantitative lipidomics show that ER membranes are dominated by phosphatidylcholine and mono-unsaturated glycerophospholipids, consistent with a highly compressible bilayer. Although proteomics suggests a functional specialization of ER tubules and an enrichment of tubule-associated proteins therein, the lipidome of an ER tubule-enriched isolate is indistinguishable from the general ER, indicating that principal ER architectures share a common lipid composition. Integration of lipidomic data with bioinformatic analyses of transmembrane helices further demonstrates that the physicochemical features of ER lipids mirror those of ER-resident membrane proteins, including reduced hydrophobicity and increased polarity compared to plasma membrane proteins. These findings support a coordinated evolution of ER proteins and lipids based on shared biophysical constraints. Together, this work provides a definitive characterization of the mammalian ER lipidome and suggest that membrane properties are maintained across the entire ER.

## Introduction

The endoplasmic reticulum (ER) is a structurally and functionally complex network of interconnected sheets and tubules. It carries out essential functions in Ca^2+^ homeostasis, membrane protein insertion, protein quality control, and lipid synthesis. Its structure covers at least two major architectures: ribosome-enriched sheets containing the protein-biogenesis machinery often found in the perinuclear region, and highly curved tubules stabilized by specialized proteins such as reticulons populating the cell periphery (Voeltz *et al*, 2006). Although these regions have been described extensively by high resolution imaging (West *et al*, 2011; Puhka *et al*, 2012; Nixon-Abell *et al*, 2016; Sawyer *et al*, 2024), and although they seem to differ in their biophysical properties (Goujon *et al*, 2019), their precise molecular composition remains poorly defined. In particular, it is unclear whether ER tubules establish a lipid environment that is distinct from the rest of the organelle.

The ER must provide a membrane environment compatible with the insertion and folding of diverse membrane proteins differing in length, side chain bulkiness, and polarity (Sharpe *et al*, 2010; Hegde & Keenan, 2021; Lorent *et al*, 2025). Membrane protein insertases have been suggested to use bilayer thinning and bilayer distortion as a common insertion mechanism (McDowell *et al*, 2020; Pleiner *et al*, 2020; Wu & Rapoport, 2021; McDowell *et al*, 2023).

The ER bilayer is soft and compressible through a high content of loosely packing lipids, in contrast to the more ordered, cholesterol-rich membranes of later secretory compartments (Holthuis & Menon, 2014; Renne & Ernst, 2023). Determining the lipid composition of the mammalian ER and its subdomains is essential for understanding how the organelle supports membrane protein insertion and folding, and how aberrant lipid metabolism contributes to lipotoxicity and ER stress (Listenberger *et al*, 2003; Volmer *et al*, 2013; Piccolis *et al*, 2019; Ernst *et al*, 2024).

A major challenge for such endeavor has been the isolation of ER membranes with sufficient purity, which is hampered by physical contact sites between the ER and other organelles (Phillips & Voeltz, 2016; Voeltz *et al*, 2024). Hence, comprehensive and quantitative analyses of the mammalian ER and its major subdomains remain limited.

Here, we introduce mammalian MemPrep, an approach that enables the isolation of ER membranes from HEK293T cells at a purity suitable for quantitative proteomics and lipidomics. As a bait for an affinity purification, we use cleavable, epitope-tagged variants of SEC61β and REEP5, which we express in HEK293T cells. SEC61β is found across the entire ER membrane network, while REEP5 is selectively enriched in ER tubules. Hence, the MemPrep isolates are representative the composition of the entire ER and the tubular ER in the case of SEC61β and REEP5, respectively.

The proteomic analysis confirms the high purity of our preparation and the expected distribution of protein machineries: curvature-stabilizing proteins such as reticulons are enriched in REEP5 isolates, whereas sheet-associated factors, including translocon components and ribosomal proteins, are significantly more enriched in SEC61β isolates. Thus, ER tubules can be selectively enriched from other ER subdomains. Remarkably, the lipid compositions of the two MemPrep isolates are nearly identical. Both feature high levels of phosphatidylcholine (PC), low cholesterol, and an enrichment of mono-unsaturated acyl chains, consistent with a relatively thin and highly compressible bilayer optimized for both membrane protein insertion and extraction. These features align with the transmembrane-helix properties of ER-resident single-pass proteins, which we find to be significantly less hydrophobic and bulkier than plasma membrane counterparts.

Together, these results provide a quantitative view on the ER membrane in mammalian cells. The lipidome of isolates from ER tubules are remarkably similar to the rest of the ER despite distinct proteomes. Hence, mammalian MemPrep offers a robust platform for dissecting ER compositions and for studying how perturbations in lipid or protein homeostasis influence ER structure and function.

## Results

We were interested in isolating ER membranes from HEK293T cells for a quantitative characterization via proteomics and lipidomics. To this end, we adapted the MemPrep procedure originally developed for the isolation of organelle membranes from *Saccaromyces cerevisiae* (*S. cerevisiae*) (Reinhard *et al*, 2023, 2024). Mammalian MemPrep relies on a gentle, detergent-free, mechanical lysis of the cells in a hypertonic buffer followed by differential centrifugation to separate ER-derived microsomes from mitochondria-derived membranes. Next, larger organelle fragments are disrupted by brief pulses of sonication, and the resulting vesicles are subjected to affinity purification using magnetic dynabead-coupled antibodies directed against the cleavable tag of the bait protein. Specifically bound, ER-derived membrane vesicles are washed with harsh, urea-containing buffers and selectively released by proteolytically cleaving the bait tag. Compared to the MemPrep procedure in yeast, we tested various means of cell disruption and optimized the differential centrifugation protocol. Apart from these modifications, the original procedures including sample handling, incubation times, buffers, antibodies and affinity matrix proved effective and were repurposed for isolating mammalian ER membranes.

As a bait for the immunoisolation of the ER, we used an N-terminally tagged variant of SEC61β, which is part of the SEC61 translocon reported to localize to ER sheets (Hartmann *et al*, 1994; Shibata *et al*, 2010; Pfeffer *et al*, 2015; Dong *et al*, 2018). However, as a fluorescently-tagged protein and depending on the degree of overexpression SEC61β has also been used as a general ER marker that localizes across the entire ER membrane network (Voeltz *et al*, 2006; Zurek *et al*, 2011; Nixon-Abell *et al*, 2016). For isolating membrane vesicles derived from ER tubules, we used a C-terminally tagged variant of Receptor expression-enhancing protein 5 (REEP5; DP1) localizing to ER tubules and stabilizing membrane curvature (Voeltz *et al*, 2006; Hu *et al*, 2008; Shibata *et al*, 2010; Kontou *et al*, 2022) (**Fig. 1A**). When attached to the N-terminus of SEC61β, the bait tag consistent of a 3xFLAG tag, a 3C protease cleavage site, and a MYC epitope, while the order was reversed in the C-terminal tag of REEP5 (**Fig. 1A**). Successful retroviral transduction of HEK293T cells was validated by immunoblotting using antibodies directed either against REEP5, SEC61β or the FLAG epitope, using anti-GAPDH immunoblots to validate equal loading (**Fig. 1B, C**). The REEP5 antibody (**Fig. 1B**) and the SEC61β antibody (**Fig. 1C**) detected both the endogenous and tagged forms of REEP5 and SEC61β, respectively. This allowed us to estimate a ≈2-fold overabundance of REEP5 in the stable cell line, as the level of the tagged variant was found at a similar level as the endogenous protein (**Fig. 1B**). The tagged variant of SEC61β was expressed at a lower level than its endogenous counterpart (**Fig. 1C**). Using anti-FLAG immunoblots, we validated the presence of the affinity tag in either SEC61β- or the REEP5-bait expressing cells (**Fig. 1B, C**). These experiments provided evidence for a similar expression level of the respective bait constructs in the two stable cell lines (**Fig. 1B**). We decided to go one step further and compared the proteomes of wildtype HEK293T cells with the two cell lines using TMT-multiplexed, untargeted protein mass spectrometry (**Suppl. Fig. S1A, B**). This experiment revealed that bait proteins have only a minimal, neglectable impact on the cellular proteome (**Suppl. Fig. S1A, B**). We did not find evidence for a systematic deregulation of proteins known to localize exclusively to ER tubules or other ER subdomains. Furthermore, quantitative proteomics validated the results from immunoblotting (**Fig. 1B, C)**: Expression of bait-SEC61β has barely any impact on the total cellular level of SEC61β (**Suppl. Fig. S1A**) while the expression of the REEP5-bait results in a 1.8-fold overabundance of REEP5 (**Suppl. Fig. S1B**).

**Figure 1:**
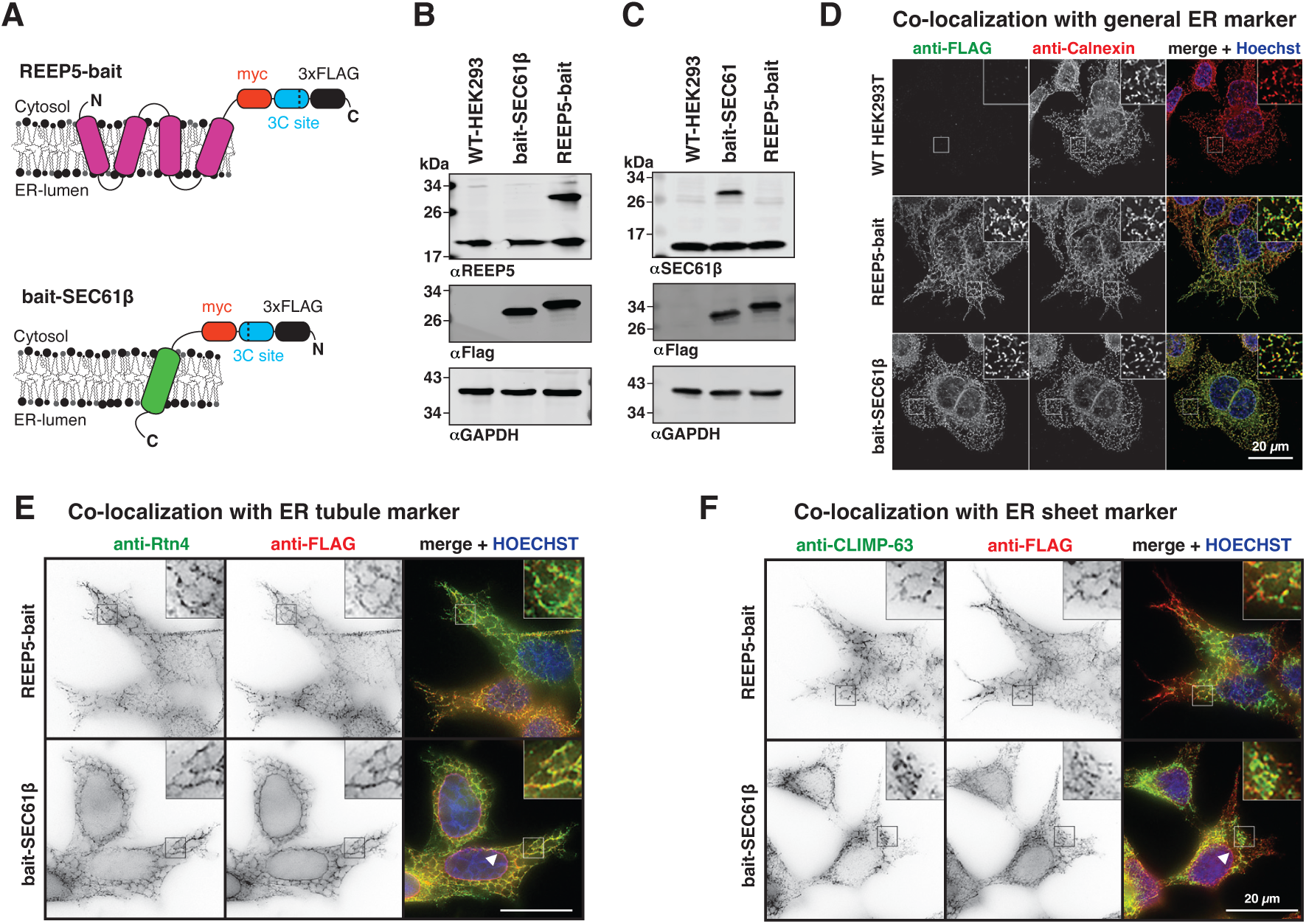
Expression of proteins in the ER of HEK293T cells for MemPrep. **A:** Schematic representation of the bait-SEC61β and REEP5-bait proteins and their membrane topology. **B:** Immunoblot analysis of cell lysates from HEK293T WT, and bait-SEC61β or REEP5-bait expressing cells using monoclonal antibodies directed against REEP5, the FLAG-tag, or GAPDH as a loading control. Note that the αREEP5 antibody detects both endogenous and tagged REEP5. **C**: Immunoblot analysis as in (B) using a polyclonal antibody directed against SEC61β. **D:** Immunofluorescence microscopy analysis of HEK293T WT and bait-SEC61β or REEP5-bait expressing cells after deconvolution. An antibody directed against Calnexin was used as an ER marker, while the bait proteins were detected using a monoclonal αFLAG antibody. **E:** Immunofluorescence analysis of fixed cells using an αRtn4 antibody as a marker for ER tubules, Hoechst33342 to stain nuclei, and αFLAG antibodies to detect either bait-SEC61β or the REEP5-bait without deconvolution. **F**: Immunofluorescence as in (E) using αCLIMP63 as a marker for ER sheets. White arrowheads mark the nuclear envelope. Micrographs are representatives of n = 3 biological replicates.

Next, we wanted to confirm normal ER localization of tagged SEC61β and REEP5 variants via immunofluorescence with antibodies directed against the FLAG epitope and against Calnexin as an ER membrane marker (**Fig. 1D**). Expectedly, we detected FLAG-positive signals only in cells with a bait construct (**Fig. 1D**) and robust colocalization of the FLAG-tagged proteins with the ER-marker Calnexin (**Fig. 1D**). Hence, the REEP5-bait and bait-SEC61β localize to the ER. We found no evidence that the expression of the bait constructs disrupts the tubule-to-sheet ratio or other aspects of the ER architecture, but distinguishing ER sheets and ER tubules is challenging in HEK293T cells.

To study localization of the bait proteins within the ER membrane network, we performed colocalization analysis of the bait proteins with either RTN4 as a tubular marker (**Fig. 1E**) (Voeltz *et al*, 2006) or CLIMP63 as a marker for ER sheets (Klopfenstein *et al*, 2001; Shibata *et al*, 2010). The REEP5-bait protein featured a near perfect colocalization with RTN4 and was found enriched in the peripheral, tubular ER (**Fig. 1E**). Bait-SEC61β was found throughout the entire ER membrane network including the nuclear envelope (**Fig. 1E**, white arrowhead), which was not populated by RTN4. The colocalization of ER sheet marker CLIMP63 with the REEP5-bait was significant, but weak in the peripheral regions of the ER (**Fig. 1F**). CLIMP63 was not found in the nuclear envelope (**Fig. 1F**, white arrowhead) but colocalized with bait-SEC61β in other regions of the ER, in particular the perinuclear ER membrane network (**Fig. 1F**). Manders’ M1 coefficient analysis (colocalization of the bait protein with an ER subdomain marker) revealed higher values for the REEP5-bait with RTN4 than with CLIMP63 consistent with a preference of the REEP5-bait for ER tubules (**Suppl. Fig. S1C**). Surprisingly, also bait-SEC61β showed preference for ER tubules albeit less pronounced than the REEP5-bait and with a wider spread of the Manders’ coefficient (**Suppl. Fig. S1D**). This is consistent with a rather promiscuous localization of bait-SEC61β across the entire ER membrane including the nuclear envelope. We like to stress, that the thresholded Mander’s coefficient, unlike the Pearson coefficient, is independent on the signal intensity and that the extent of colocalization is dependent on the resolution of the imaging modality. This is particularly relevant for imaging the ER, because dense tubular networks in the cell periphery can appear like ER sheets in diffraction-limited microscopy (Nixon-Abell *et al*, 2016). Furthermore, the edges of ER sheets are populated by curvature-stabilizing proteins also found in ER tubules (Shibata *et al*, 2010; Shemesh *et al*, 2014), and ER sheets show different degrees of fenestration dependent on the cell type and the cell cycle phase (Puhka *et al*, 2007, 2012; Nixon-Abell *et al*, 2016). Consistent with our microscopic data (**Fig. 1E, F**) and because ER sheets may be biochemically inseparable from ER tubules, we use SEC61β as a general ER marker.

### Mammalian MemPrep isolates and enriches ER membranes

Mild cell lysis was achieved by gentle and Dounce homogenization. The lysates were subjected to differential centrifugation to enrich for ER membranes (**Fig. 2A**). A first centrifugation (1,000 x g, 10 min, 4°C) removed nuclei and unbroken cells in the pellet. The resulting supernatant was centrifuged (10,000 x g, 10 min, 4°C) to remove mitochondria, which can represent a major contaminant when isolating ER membranes (Vance, 1990; Leiro *et al*, 2023). To yield a crude microsome preparation enriched in ER membranes, the supernatant was centrifuged (100,000 x g, 60 min, 4°C) to pellet crude microsomes and to discard cytosolic proteins in the pellet.

**Figure 2:**
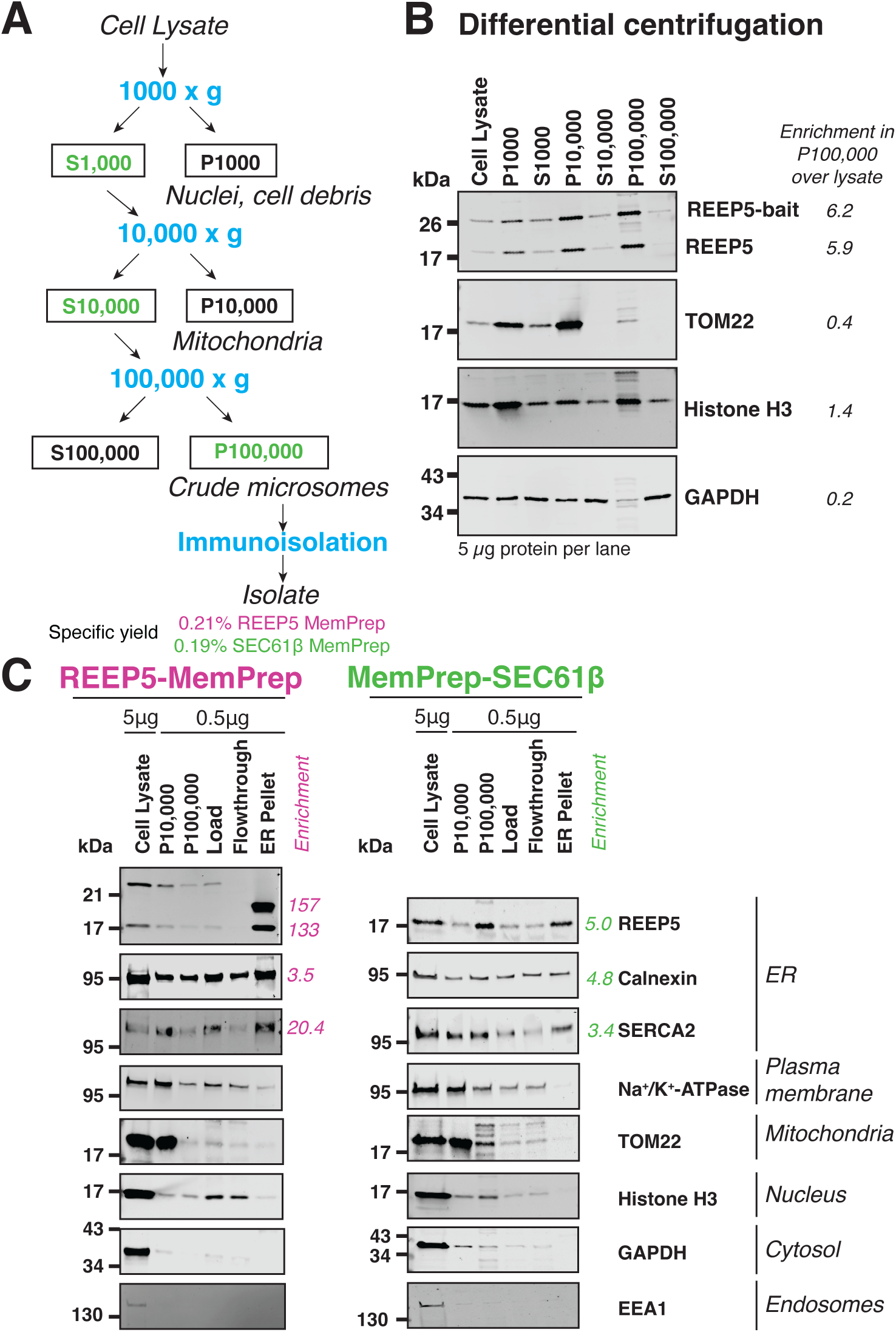
Mammalian MemPrep yields highly enriched ER preparations. **A:** Schematic representation of the mammalian MemPrep workflow. **B:** Immunoblotting using antibodies directed against REEP5, TOM22, Histone H3, and GAPDH were used to validate the differential centrifugation procedure. TOM22, Histone H3, and GAPDH are markers for mitochondria, the nucleoplasm, and the cytosol, respectively. 5 µg of protein from the indicated fraction were separated by SDS-PAGE and subjected to immunoblotting. The relative enrichment and depletion of organelle-specific markers was calculated by densitometry. **C:** Indicated fractions and protein amounts from the MemPrep procedure via the REEP5-bait or bait-SEC61β were subjected to SDS-PAGE and immunoblotting using antibodies against the indicated organelle marker proteins. The relative enrichment of ER markers in the final isolates was determined by densitometry.

We validated the differential centrifugation procedure by immunoblotting using different subcellular markers (exemplary shown for samples from REEP5-bait expressing cells) (**Fig. 2B**). To document the enrichment of ER proteins (rather than the yield) in each fraction, we loaded equal protein amounts (5 µg). Both the endogenous and tagged form of REEP5 were enriched in the P10,000 and P100,000 membrane fractions (**Fig. 2B**). This was expected, because abundant, cytosolic proteins are removed in the final step of the differential centrifugation procedure. Immunoblotting using antibodies directed against the outer mitochondrial membrane protein TOM22 revealed a partial removal of mitochondrial membranes by a centrifugation at 1,000 x g, and a near-complete removal by centrifugation at 10,000 x g (**Fig. 2B**). Hence, the crude microsomal P100,000 fraction that is subjected to immunoisolation is already significantly depleted of mitochondrial contaminations. Immunoblotting against Histone H3 revealed an enrichment in the P1,000 pellet fraction, where unbroken cells and intact nuclei are expected (**Fig. 2B**). The continuous presence of Histone H3 in other fractions of the procedure suggests that some nuclei are disrupted during cell lysis thereby releasing their content (**Fig. 2B**). Immunoblotting using antibodies directed against GAPDH revealed a significant depletion of GAPDH in the crude microsomal fraction (**Fig. 2B**).

Overall, we conclude that differential centrifugation enriches for ER membranes (**Fig. 2B**). This is supported by a quantitative analysis of the immunoblots, suggesting a 5.9- and 6.2-fold enrichment of the endogenous and the tagged REEP5 variant, respectively, in the P100,000 fraction. Yet this enrichment prior to the immunoisolation comes at a significant price, as ≈90% of ER membranes are lost during differential centrifugation (9.2% and 10.1% yield for the MemPrep via REEP5 and SEC61β, respectively). Intriguingly, the total protein yield was even lower (**Suppl. Table 1**), presumably because soluble proteins remain in the supernatant and discarded.

Next, we subjected the crude microsomal preparation to brief pulses of sonication to separate large clumps of aggregated vesicles and to disrupt larger ER fragments into smaller vesicles as previously established for isolating microsomes and more recent immunoisolation studies (Dallman *et al*, 1969; Stan *et al*, 1997; Reinhard *et al*, 2023, 2024). Even though this procedure induces transient membrane ruptures that result in a partial release of ER luminal content, there is no indication that sonication causes membrane mixing through fusion of ER-derived microsomes with non-ER-derived vesicles (Reinhard *et al*, 2024).

Next, the crude microsomal preparation was subjected to an immunoisolation procedure using magnetic protein G dynabeads sparsely decorated with anti-FLAG antibodies. After incubation of the ER-enriched membrane preparation with the affinity matrix and extensive washing with buffers containing 0.6 M urea, we eluted the specifically bound vesicles by cleaving the bait tag with affinity-purified GST-HRV3C protease. The overall MemPrep procedure was validated for both isolations via the REEP5-bait (**Fig. 2C**) and bait-SEC61β (**Fig. 2D**). Note that the electrophoretic mobility of the REEP5-bait construct changes upon cleaving the bait tag (**Fig. 2C**). While REEP5, Calnexin and the ER-localized Ca^2+^ pump SERCA2 were robustly detected and enriched in the final isolate (**Fig. 2C, D**), the absence of the Na^+^/K^+^-ATPase (plasma membrane), TOM22 (mitochondrial outer membrane), Histone H3 (nucleus), EEA1 (endosomes), and GAPDH (cytosol) demonstrated a significant depletion of other cellular compartments in the isolate (**Fig. 2C, D**). These findings demonstrate a significant enrichment of ER membranes through mammalian MemPrep. As expected for the preparation of an organelle membrane, the total protein yield was low (**Supplementary Table 1**).

### Proteomics reveals ER membrane isolation via mammalian MemPrep

While the success of immunoisolation is often documented by the loss of several organelle-specific markers, we decided to go one step further. For a more rigorous validation, we measured the enrichment of known ER proteins over the lysate by TMT-multiplexed, untargeted protein mass spectrometry. In the case of the REEP5 MemPrep, we detected 6060 proteins including 437 ER-annotated proteins (**Fig. 3A**). In the case of the SEC61β MemPrep we quantified the relative abundance of 5191 proteins including 365 ER-annotated proteins (**Fig. 3B**). Volcano plots revealed the specific enrichment and depletion of proteins in mammalian MemPreps over the respective cell lysates (**Fig. 3A, B**). In both the REEP5 and the SEC61β MemPrep, we found most ER-annotated proteins strongly enriched in the isolate and more so than proteins annotated to other organelles (**Fig. 3A-D**). Overall, the immunoisolations via the REEP5-bait yielded higher enrichments of ER-annotated proteins than observed for bait-SEC61β (**Fig. 3C, D**). This suggests a higher purity of the REEP5 MemPrep.

**Figure 3:**
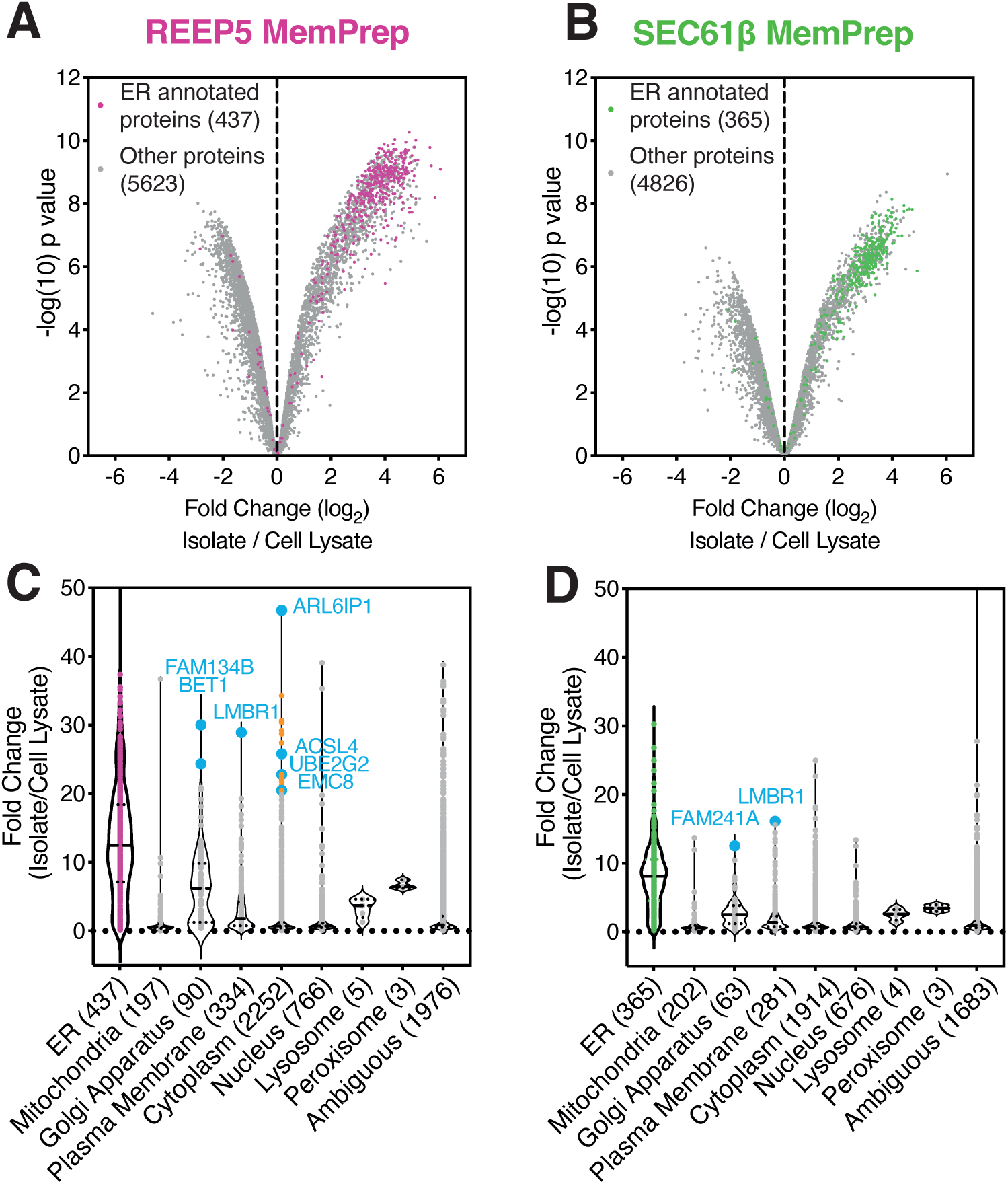
Quantitative proteomics reveals a specific enrichment of ER proteins by mammalian MemPrep. **A:** Limma analysis after non-targeted, TMT-labeling proteomics consistently detected 6060 proteins in the cell lysate and the REEP5 MemPrep isolate. Proteins annotated and categorized as ER-proteins (purple) are enriched in the final isolate compared to the cell isolate. **B:** Limma analysis after non-targeted, TMT-labeling proteomics consistently detected 5191 proteins in the cell lysate, and in the SEC61β MemPrep. ER-annotated proteins (green) are prominently enriched in the final isolate. **C, D:** Binning the limma fold change according to a GO-term analysis ‘subcellular localization’ indicates the limma fold change enrichment of proteins from different organelles in the isolate over the whole cell lysate for the REEP5 MemPrep (C) and the SEC61β MemPrep (D). Proteins highlighted in blue are examples for known ER proteins misannotated or miscategorized as non-ER proteins. Likewise, proteins highlighted in tangerine are >20-fold enriched proteins in the REEP5 MemPrep, classified as cytoplasmic but indeed associated with the ER membrane. Data information: The average from n = 3 biological replicates in (A, B) is represented as a single datapoint.

Even though mammalian MemPrep was optimized to provide insight into subdomain specific composition by comparing ER tubules with the rest of the ER, quantitative proteomics revealed that the two isolates via the REEP5-bait and bait-SEC61β were largely correlated when considering either all 4814 proteins consistently detected in the triplicates of both isolations (**Suppl. Fig. S2A**) or only the 357 ER-annotated proteins (**Suppl. Fig. S2B**). Nevertheless, the overall similarity of both isolates is not entirely surprising as they both isolate membrane fragments from a continuous membrane network.

Next, we systematically retrieved the GO term ‘Cellular component’ for each detected protein and binned them into the categories ‘ER’, ‘Mitochondria’, ‘Golgi apparatus’, ‘Cytoplasm’, ‘Nucleus’, ‘Lysosome’, or ‘Peroxisome’ (**Fig. 3C, D**) based on the first GO term retrieved from Uniprot (Huntley *et al*, 2014). If the retrieved GO term was distinct from the above-mentioned categories or had multiple of them, the protein subcellular localization was categorized as ‘Ambiguous’ (**Fig. 3C, D**).

Plotting the enrichment of proteins sorted by subcellular localization revealed higher enrichments of ER-annotated proteins over proteins primarily annotated to other organelles such as mitochondria, the Golgi apparatus, or the nucleus (**Fig. 3C, D**). Finding non-ER proteins in an ER proteome is not surprising, because a very large number of proteins are first delivered to the ER, before they are sent to other cellular destinations. Nevertheless, manually inspecting cases of highly enriched proteins revealed not annotated to the ER revealed either misannotations or misinterpretations from binning the available information. For example, the two Golgi-annotated proteins with the highest enrichment in the REEP5 MemPrep are known to reside in the ER membrane: the reticulophagy receptor FAM134B (Khaminets *et al*, 2015) and the SNARE-protein BET1 involved in ER-to-Golgi transport (Newman *et al*, 1990; Lian & Ferro-Novick, 1993) (**Fig. 3C**). Likewise, all 23 highly enriched proteins annotated as ‘cytoplasmic’ in the REEP5 MemPrep are indeed known ER proteins or at least predicted to reside in the ER (**Fig. 3C**). These include the ADP-ribosylation factor-like protein 6-interacting protein 1 (ARL6IP1) involved in shaping the ER along its interacting Inositol polyphosphate 5-phosphatase K (Yamamoto *et al*, 2014; Dong *et al*, 2018), the Acyl-CoA Synthetase Long-Chain Family Member 4 (ACSL4) crucial for incorporating polyunsaturated fatty acids into membrane lipids (Küch *et al*, 2014), the ubiquitin-conjugating protein UBE2G2 involved in ER-associated degradation (Klemm *et al*, 2011; Tsai *et al*, 2022), and EMC8 involved in membrane protein insertion (Guna *et al*, 2017). Similar observations can be made with the SEC61β MemPrep: The highly enriched FAM241A annotated as Golgi apparatus protein has recently been identified as an ER protein (Kobayashi *et al*, 2022) (**Fig. 3D**). Despite being annotated as plasma membrane protein, Limb region 1 protein homolog (LMBR1) was highly enriched by both the REEP5 and the SEC61β MemPreps (**Fig. 3C, D**) consistent with its recently established ER localization (Choi *et al*, 2019). Thus, mammalian MemPrep provides highly enriched ER membrane preparations. The combination of MemPrep with quantitative proteomics bears the potential to even predict the localization of a protein to the ER membrane.

### TMHs in the ER are less hydrophobic and bulkier than in the plasma membrane

We were interested if the TMHs of ER membrane proteins differ from the TMHs found in other organelles along the secretory pathway such as the Golgi apparatus or the plasma membrane. We extracted the TMH sequence of 82 robustly detected, ER-annotated single-pass membrane proteins and compared their overall properties with the previously characterized 90 Golgi apparatus TMHs and 273 plasma membrane TMHs (Sharpe *et al*, 2010) using HelixHarbor, a novel, custom-built bioinformatic tool we developed for this study (**Fig. 4A-D**). The analysis could have been extended to all ER-, Golgi apparatus-, and PM- annotated proteins, but we focused on those ER proteins detected in our experiments. Consistent with previous findings (Sharpe *et al*, 2010), we found that the TMHs of ER and Golgi apparatus proteins are shorter than the TMHs in the plasma membrane (**Suppl. Fig. S3A**). These observations from bioinformatic analyses have recently been experimentally underscored by cryo-EM tomography analyzing the thicknesses of organelle membranes (Glushkova *et al*, 2026).

**Figure 4:**
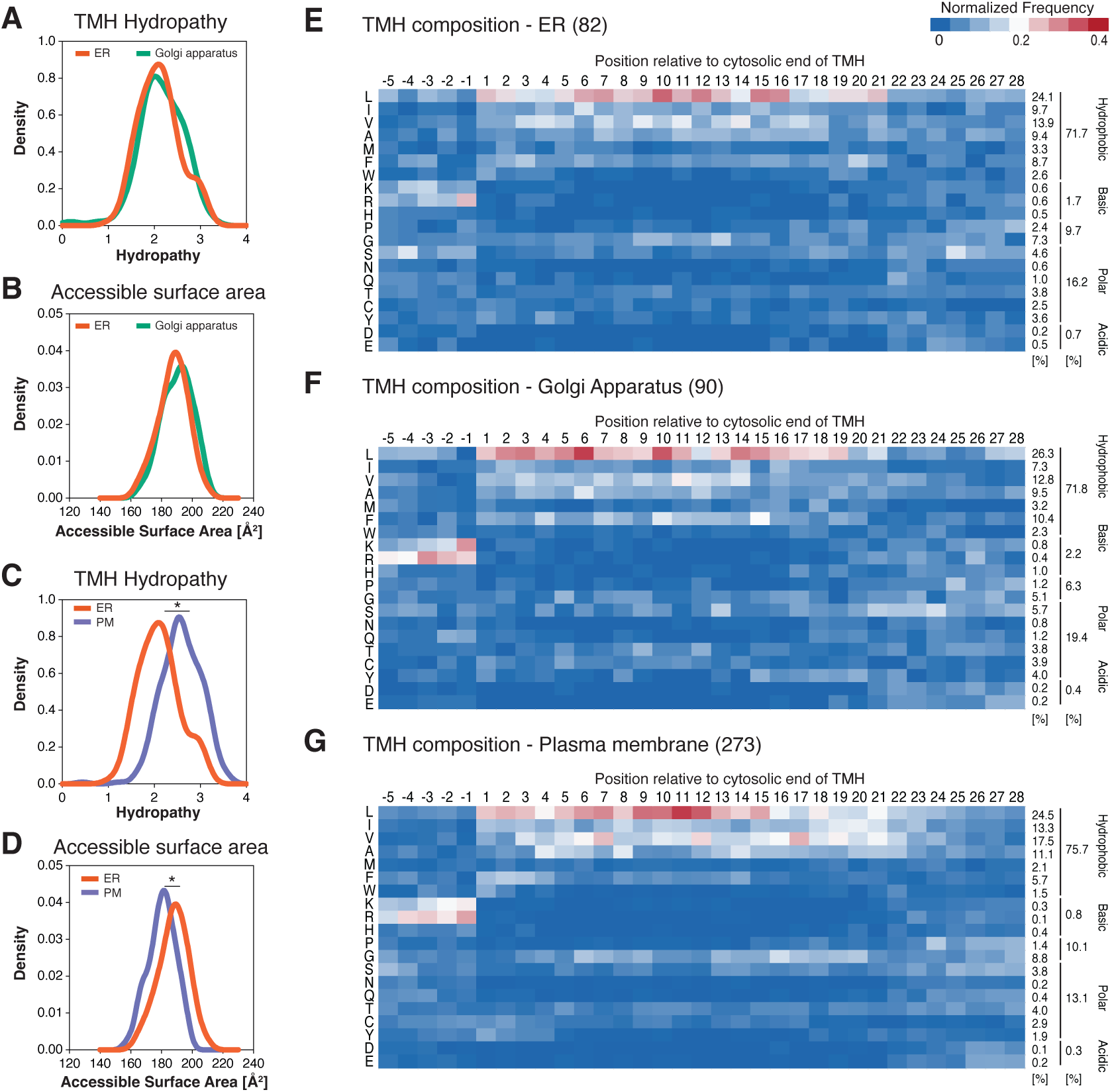
ER transmembrane domains are more polar than those in the plasma membrane. **A, B:** The predicted transmembrane helices of 82 single-pass, ER-annotated transmembrane proteins from our proteomic experiments were subjected to a statistical analysis and compared with single-pass transmembrane helices from the Golgi apparatus (90) and the plasma membrane (273) (Sharpe *et al*, 2010). The frequency of different hydropathies (A) and the average accessible surface area (B) are plotted comparing transmembrane helices from the ER (red) and the Golgi apparatus (green). **C, D:** Comparison of transmembrane helices in the ER (red) and the plasma membrane (violet) with respect to hydropathy (C) and accessible surface area (D). **E, F, G:** The relative frequencies of individual amino acids in and around transmembrane helices of single-pass proteins are plotted for the ER (E), the Golgi apparatus (F), and the plasma membrane (G). The transmembrane helices are oriented with the positively charged, juxta-membrane section facing left. Position 1 indicates the beginning of the predicted transmembrane helix. The data are normalized such that the frequencies of amino acids add up in each column to 1. The relative frequency of a specific amino acids and amino acid categories in the set of TMHs is provided on the right side of the heatmap. Statistical test: Kolmogorov-Smirnov test, T-test and Wilcoxon test were performed (A-D) each comparing two datasets. No significant differences were observed in (A) and (B). * represents qualitative significance using all three types of test in (C) and (D).

Analysis of the hydropathy (Kyte & Doolittle, 1982) and accessible surface area (Tien *et al*, 2013) revealed no significant differences between the TMHs from the ER and the Golgi apparatus (**Fig. 4A, B**), but highly significant differences between ER and plasma membrane proteins (**Fig. 4C, D**). Thus, single-pass proteins in the ER are less hydrophobic (**Fig. 4C**) and bulkier (**Fig. 4D**) than their plasma membrane counterparts, which is consistent with the number of non-polar (**Suppl. Fig. S3B**) and polar TMH residues (**Suppl. Fig. S3C**).

Next, we extended our analysis of TMH properties to a single-amino acid level and calculated the relative frequency of specific amino acids relative to the more positively charged end of the predicted TMH, which is distinct from a strict N-to-C-terminus directionality (**Fig. 4E-G**). We decided for this approach to accommodate the positive-inside-rule, which describes the general trend that single-pass TMHs are inserted into the membrane with their positive end facing the cytosol (von Heijne & Gavel, 1988; Heijne, 1992). Consequently, we studied the frequency of amino acid residues from the cytosol towards the organelle lumen (or the extracellular space, respectively) thereby following the strategy used in the pioneering work by Sharpe *et al*. (Sharpe *et al*, 2010). Again, we focused on 82 single-pass, ER-annotated proteins and compared their TMHs to previously analyzed single-pass TMHs from the Golgi apparatus or the plasma membrane (Sharpe *et al*, 2010). Reflecting the positive-inside-rule, we observed an increased frequency of positively charged residues adjacent to one end of the TMH for ER (**Fig. 4E**), Golgi apparatus (**Fig. 4F**), and plasma membrane proteins (**Fig. 4G**). Intriguingly, the cumulative frequency of positive residues in this region was higher for Golgi and plasma membrane proteins compared to the ER consistent with a recent bioinformatic study (Lorent *et al*, 2025). It is tempting to speculate that the frequency of positive residues in the juxtamembrane region is related to the negative surface charge provided by lipids on the cytosolic side of the membrane. The negative surface charge density from lipids is generally assumed to be low in the ER, but much higher in organelles of the late secretory pathway (Bigay & Antonny, 2012; Holthuis & Menon, 2014).

Confirming our expectations based on the predicted TMH length (**Suppl. Fig. S3A**), we observed a gradual decline in the relative frequency of hydrophobic and aromatic resides at about 21 amino acids for ER (**Fig. 4E**) and Golgi-associated TMHs (**Fig. 4F**). Such decline was more clearly defined for plasma membrane TMHs but only after 24 aa or more (**Fig. 4G**). Hence, the TMHs in the early secretory pathway are shorter than in the plasma membrane, which is also reflected in the number of polar and aploar residues in the TMHs (**Suppl. Fig. S3B, C**). Consistent with previous reports (Sharpe *et al*, 2010; Lorent *et al*, 2017, 2020, 2025), we also observed an asymmetric distribution of large, aromatic amino acid side chains in the TMHs of plasma membrane proteins, while we found no indication for an inverted asymmetry in our smaller dataset of single-pass ER proteins (**Fig. 4E**).

Our new finding that the TMHs of ER proteins are more polar than the TMHs in the plasma membrane (**Fig. 4C**) is also reflected by the significantly different number of apolar and polar residues in the TMHs from ER-, Golgi apparatus-, and PM-derived proteins (**Suppl. S3B, C**). These observations are consistent with recent, even larger bioinformatic datasets (Lorent *et al*, 2025). We therefore challenged our finding and performed an additional analysis using this larger dataset of all annotated human single-pass TMHs (**Fig. S3D**) and compared the hydrophobicity profiles of TMHs from the ER (215), the Golgi apparatus (260), and the PM (1322) (Lorent *et al*, 2025). This analysis further substantiated our finding that the ER and the Golgi apparatus host less hydrophobic TMHs compared to the plasma membrane. Furthermore, we observed that the ER and Golgi profiles display a conical shape with hydrophobic maxima at the center of the membrane’s hydrophobic core, while the PM TMH’s possess higher hydrophobicity in the cytoplasmic part of the membrane, compared to the exoplasmic part (**Fig. S3D**).

In summary, our bioinformatic analysis confirms known characteristics of ER resident membrane proteins with respect to TMH length and accessible surface area, and established that ER and Golgi apparatus TMHs are more polar than those in the plasma membrane

### MemPrep differentially enriches ER subdomains

Quantitative proteomics revealed that the isolation of ER membranes via two distinct bait proteins yielded overall similar proteomes (**Suppl. Fig. S1C, D**). However, we also observed a pronounced enrichment of known tubular proteins such as Atlastin-2, Atlastin-3, and the Receptor expression-enhancing proteins 3, 5, and 6 (REEP3, REEP5, REEP6) in the REEP5 MemPrep (Voeltz *et al*, 2006; Rismanchi *et al*, 2008; Park *et al*, 2010), and -conversely- higher enrichments of the unfolded protein response transducer IRE1α (ERN1) (Belyy *et al*, 2022), Argonate 2 (Ago 2) (Gao *et al*, 2022), and the lipid metabolic enzymes Lysophosphatidylcholine Acyltransferase 3 (LPCAT3) (Hishikawa *et al*, 2008), 1-Acylglycerin-3-phosphat-O-Acyltransferase 1 (AGPAT1) (Adachi *et al*, 2025), and the 7-Dehydrocholesterol-Reduktase (DHCR7) (Zwerger *et al*, 2009) in the SEC61β MemPrep (**Suppl. Fig. S1C, D**).

Because the selective enrichments of proteins in either the REEP5 or SEC61β MemPrep is not caused by the expression of the bait proteins (**Suppl. Fig. S1A, B**), they are indicative that some differences between ER tubules and other ER subdomains including sheets and the nuclear envelope are preserved in MemPrep isolates (**Suppl. Fig. S1C, D**). Hence, we performed a new round of REEP5 and SEC61β MemPreps in triplicates for a direct comparison of the isolates (**Fig. 5A, B**) rather than comparing the changes in abundance relative to the respective cell lysates as performed in Figure 3. Knowing that non-ER proteins are less efficiently enriched by the MemPrep procedure than ER proteins (**Fig. 3C, D**) and that the sensitivity and comprehensiveness of mass spectrometry-based proteomics experiments are reduced with increasing sample complexity (Ting *et al*, 2011; Beck *et al*, 2011), we were hoping to gain a better insight into the distribution of low abundant and challenging to quantify proteins in the two MemPrep isolates.

**Figure 5:**
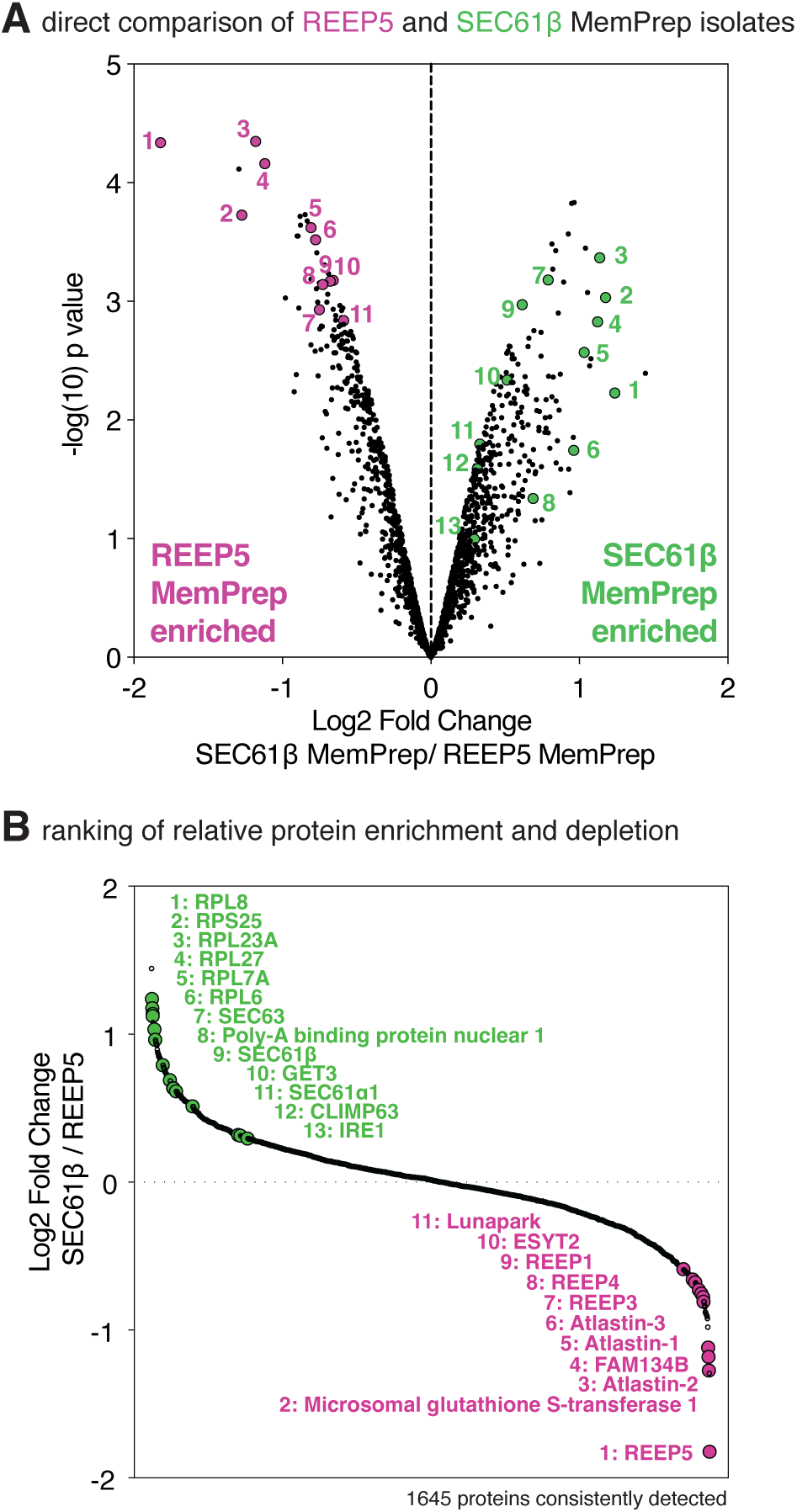
Mammalian MemPrep selectively enriches ER sheets and tubular ER membranes. **A:** Limma analysis after non-targeted, TMT-labeling proteomics on n = 3 biological replicates consistently detect 1645 proteins in mammalian MemPrep isolates via the REEP5 and the SEC61β baits. A volcano plot detects significant differences between the two isolates. Exemplary proteins known to localize to ER tubules are enriched in the REEP5 MemPrep (purple), while proteins involved in membrane protein biogenesis are enriched in the SEC61β MemPrep (green).

Indeed, we found a variety of ribosomal proteins, the Poly-A-binding protein, components of the translocon (SEC61α, SEC63), a membrane protein insertase (GET3), and the ER sheet stabilizing protein CLIMP63 (CKAP4) enriched in the SEC61β MemPrep (**Fig. 5A, B**), while the REEP5 MemPrep was selectively enriched for proteins known to reside in ER tubules including Atlastin1-3, Lunapark, Extended-Synaptotagmin (E-Synt), and REEP1, 3-5 (**Fig. 5A, B**). This suggests that mammalian MemPrep provides a preparative approach to differentially enrich ER membrane subdomains. The our proteomic analysis demonstrates that the compositional characteristics of ER subdomains are at least partially preserved during the preparation. Encouraged by these observations, we were interested if lipids, like proteins, show a preference for different ER subdomains.

### The lipid composition of the ER in HEK293T cells

Having established mammalian MemPrep, we were interested in the lipid composition of the ER membrane. We isolated ER membranes via the REEP5-bait and bait-SEC61β, extracted lipids using a two-step extraction (Ejsing *et al*, 2009), and subjected them to quantitative shotgun lipidomics. On average, only 1.2% and 0.2% of the total lipids amount in the whole cell lysate was recovered with the final isolate in the case of the REEP5 and the SEC61β MemPrep, respectively (**Supplementary table 1**).

As a control, we also tested the impact of the bait constructs on the HEK293T whole cell lipidome (**Suppl. Fig. 4A-J**). Overall, the lipid composition of the virally transduced cells was indistinguishable from HEK293T cells with only minor impact on the level of CL and lysolipids (**Suppl. Fig. 4A-J**).

The lipid compositions of the two MemPrep isolates differed substantially from the composition of the respective whole cells (**Fig. 6A-H; Suppl. Fig. S4K, L**). Compared to each other, the isolates from the REEP5 and the SEC61β MemPrep featured identical lipidomes (**Fig. 6A-H; Suppl. Fig. S4K, L**) despite their different proteomes enriched either in proteins with preference for ER tubules or ER sheets, respectively (**Fig. 5**).

**Figure 6:**
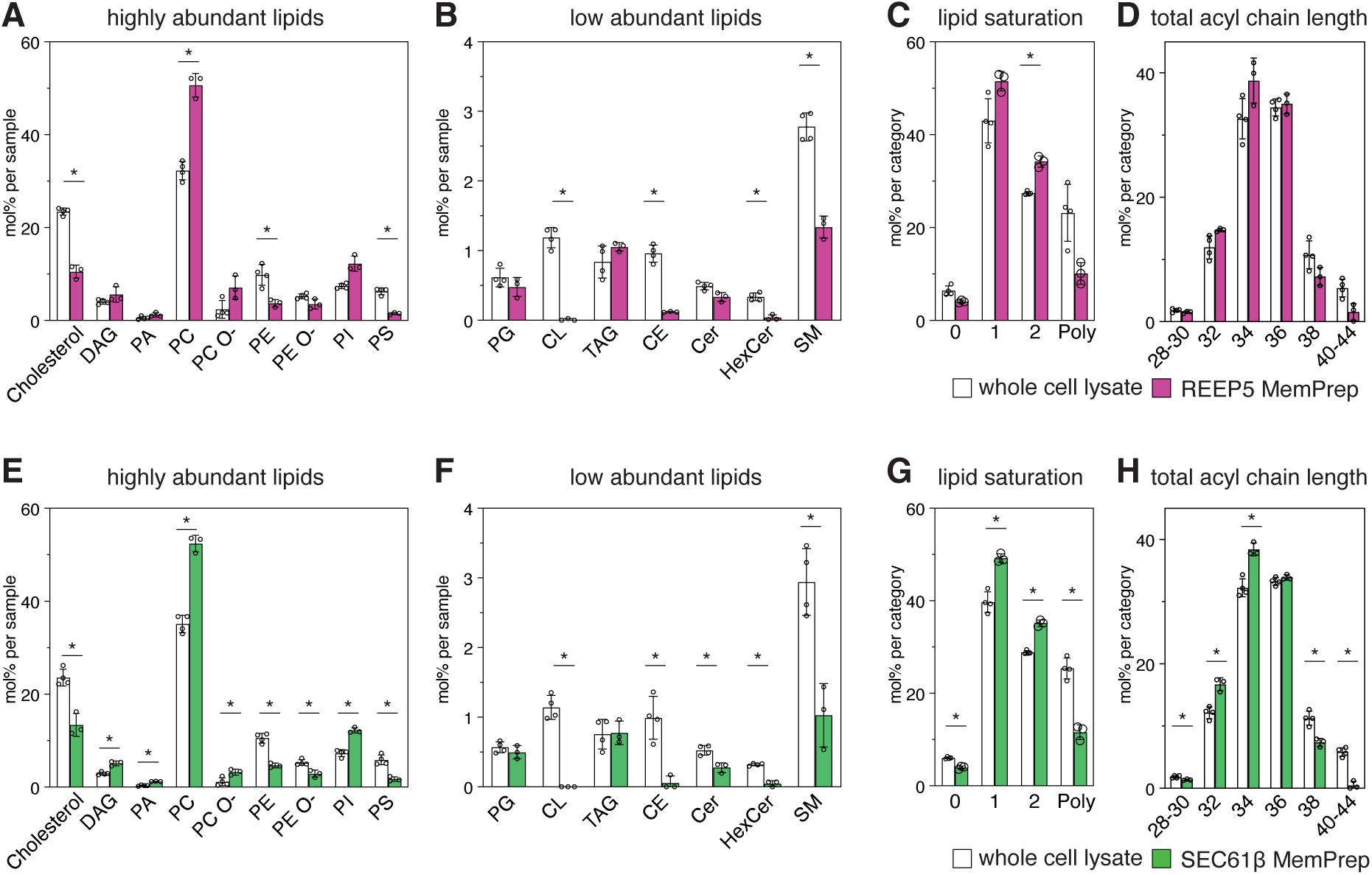
Mammalian MemPrep established the lipid composition of the HEK293T cell ER and its enriched subdomains. **A:** Quantitative lipidomics reveals the lipid class composition given as mol% of all identified lipids in the REEP5-MemPrep isolate and compared with the corresponding whole cell lysate. Except for cholesterol, each bar corresponds to a lipid class composed of several lipid species (*highly abundant lipids* - DAG: diacylglycerol, PA: phosphatidic acid, PC: phosphatidylcholine, PC O-: Phosphatidylcholine (-ether), PE: phosphatidylethanolamine, PC O-: Phosphatidylethanolamine (-ether), PI: phosphatidylinositol, PS phosphatidylserine). **B:** Lipid class composition of low abundant membrane glycerophospholipids except for lysolipids are given as mol% of all consistently identified lipids in the sample (PG: phosphatidylglycerol, CL: cardiolipin, TAG: Triacylglycerol, CE: Cholesterol-ester, CE: Ceramide, HexCer: Hexosylceramide, SM. Sphingomyelin). **C:** Lipid unsaturation of whole cell lysates and the resulting REEP5 MemPrep isolate is given as the total number of double bonds in membrane glycerolipids except for CL (i.e. DAG, PA, PC, PC O-, PE, PE O-, PI, PS, PG) as mol% in this group. All lipids with more than two unsaturations in the lipid acyl chains are categorized as Poly-unsaturated lipids (Poly). **D:** The relative abundance of membrane glycerolipids with different total acyl chain lengths (calculated as the sum of both fatty acyl chains) in whole cell lysates and REEP5 MemPrep isolates is plotted in mol% of this category. **E-H:** Identical analysis as in (A-D) comparing the lipidome SEC61β MemPrep isolates with the corresponding whole cell lysate. Data information: Data from n = 4 biological replicates in (A-H) are represented as individual data points and as the mean ± SD. Statistical test: Multiple unpaired T-tests with Welch’s correction were performed by the Holm-Šídák method with a P-value threshold of 0.05. * represents qualitative significance.

Even though the contribution of different ER subdomains to these isolates remains unclear, our data suggest that ER tubules and ER sheets feature similar or -at least-not grossly different lipid compositions. Consistent with previous reports (Radhakrishnan *et al*, 2008; van Meer *et al*, 2008; Vance, 2015; Harayama & Riezman, 2018; Sokoya *et al*, 2022; Foged *et al*, 2025), we found that PC is the most abundant lipid class in the ER with a cumulative abundance of >50 mol%, while the cholesterol levels are low (≈12 mol%) (**Fig. 6A, E**). Compared to the whole cell lipidome, the ER membranes featured dramatically lower levels of phosphatidylserine (PS) (**Fig. 6A, E**), cardiolipin (CL) and cholesterol-esters (CE) (**Fig. 6B, F**). These findings underscore the purity of our preparation, as PS lipids are enriched in endosomes and the plasma membrane, cardiolipin (CL) in the inner mitochondrial membrane, and cholesterol-esters (CE) in lipid droplets (Leventis & Grinstein, 2010; Harayama & Riezman, 2018; Foged *et al*, 2025). Ether PC (PC O-) and ether PE (PE O-) lipids contain ether-linked fatty alcohols in the *sn-1* position. Intriguingly, we find PC O- enriched in ER membranes, while PE O- is depleted (**Fig. 6A, E**). These observations are consistent with the notion that ER-synthesized etherlipids disperse broadly to establish distant membrane territories (Foged *et al*, 2025).

Compared to the whole cell lysate, we also detected reduced levels of sphingomyelin (SM) and Hexosylceramide (HexCer) in isolated ER membranes (**Fig 6B, F**). This is consistent with the biosynthetic pathway of sphingolipids such as SM and HexCer: Ceramide, which is synthesized in the ER, first must reach different subdomains of the Golgi apparatus to serve as a substrate for the biosynthesis of SM and HexCer (D’Angelo *et al*, 2013). Hence, it can be expected that complex sphingolipids are found only at reduced levels in the ER. It is intriguing, though, that the depletion of HexCer in ER membranes is more complete than that of SM (**Fig. 6B, F**). We speculate that a retrograde flux of SM to the ER via vesicular traffic is possible, while Glucan-containing HexCer may be selectively sequestered away from such retrograde carriers by their interaction with lectins and the juxtamembrane region of transmembrane proteins.

Quantitative lipidomics provided also a deeper insight into the composition of the lipid acyl chains (**Fig. 6C-D, G-H**). We found a robust enrichment of glycerophospholipids with one or two monounsaturated fatty acyl chains in the ER at the expense of lipids with three or more double bonds in the acyl chain region (**Fig. 6C, G**). This underscores the existence of a lipid gradient along the secretory pathway, where monounsaturated fatty acyl chains are enriched in the early secretory pathway to reduce membrane compressibility and increase membrane fluidity (Schneiter *et al*, 1999; Bigay & Antonny, 2012; Holthuis & Menon, 2014; Renne & Ernst, 2023; Reinhard *et al*, 2024; Foged *et al*, 2025), while asymmetric lipids with one saturated and one polyunsaturated fatty acyl chain are enriched in organelles of the late secretory pathway including the plasma membrane (Keenan & Morré, 1970; Antonny *et al*, 2015; Manni *et al*, 2018). These changes of the lipid acyl chains are associated with biophysical changes of the membrane properties along the secretory pathway as observed by molecular probes reporting on lipid packing and membrane tension (Goujon *et al*, 2019; López-Andarias *et al*, 2021, 2022; Wong & Budin, 2024). We also found evidence for differences in the acyl chain length of the glycerophospholipids (**Fig. 6D, H**). The abundance of lipids with short fatty acyl chains was higher in the ER, the abundance of lipids with long acyl chains was reduced. Hence, the ER is enriched for lipids with short, monounsaturated acyl chains.

Next, we took a closer look at different species of Ceramide (Cer) and SM, which are found at a considerable level in the ER in contrast to HexCer (**Fig. 4B, F)**. Intriguingly, ceramides enriched in the ER feature more hydroxylations than Cer in the whole cell lysate irrespectively of the bait protein used for ER isolation (**Suppl. Fig. S5A-D**). This characteristic hydroxylation was not observed for SM, whose species distribution is indistinguishable for the ER and whole cell lysates (**Suppl. Fig. S5A-D**). The increased polarity of Cer in the ER is intriguing, as it is reminiscent of the increased polarity of single-pass membrane proteins in the ER (**Fig. 4A, C**). Future work will be required to dissect a possible role of Cer hydroxylation for ER properties and functions.

Taken together, we have established the lipid composition of the ER in HEK293T cells. MemPrep isolates enriching different subdomains of the ER (**Fig. 5**) feature almost identical lipidomes (**Fig. 6**). While we cannot exclude the possibility that some of the sub-organelle organization is lost during the mild mechanical cell lysis under hypertonic conditions, these quantitative data suggest that ER tubules have a similar lipid composition as the rest of the ER. There is an urgent need for new, complementary technologies to directly image the distribution of lipids in different ER subdomains to further validate this interpretation (Iglesias-Artola *et al*, 2025; Böhlig *et al*, 2025).

## Discussion

We have established mammalian MemPrep as a technology to isolate ER membranes for a quantitative analysis of their composition. The entire procedure takes approximately 16 h from harvesting the cells to freezing the isolate. While mammalian MemPrep is slow compared to similar, complementary approaches of organelle isolations (Sokoya *et al*, 2022; Fasimoye *et al*, 2023; Abu-Remaileh *et al*, 2017; Hundley *et al*, 2024; Foged *et al*, 2025), it provides utmost purity and insight into the composition of the ER and ER tubules: 1) Differential centrifugation combined with immunoisolation maximizes the purity of the final isolate. 2) Cleavable tags allow for the specific capture and selective elution of ER membrane vesicles, thereby further enhancing the purity of the isolate and maximizing its usability for downstream analytic procedures. 3) Using bait proteins localizing to different region of the ER, provides insight into the lipid and protein composition of ER subdomains. There are, however, also limitations and technical challenges to mammalian MemPrep: 1) We use bait constructs in virally transduced cells to differentially enrich membranes from different ER subdomains. Even though the additional expression of the bait constructs has no obvious impact on the ER membrane network (**Fig. 1C**), the cellular proteome (**Suppl. Fig. S1A, B**), and the cellular lipid composition (apart from minor changes in the levels of lysolipids and CL) (**Suppl. Fig. S4**), we cannot exclude that the mild overexpression has some impact on the structure or function of the ER. 2) Mammalian MemPrep was optimized for purity, but not for high yields. We estimate that only a small fraction (≈0.2%) of ER membranes in the cell lysate are recovered in the final isolate. 3) Mild cell disruption by Dounce homogenization in a hypertonic buffer is crucial for cracking cells open, but these procedures can disrupt normal ER architecture and might facilitate the undesired mixing of previously well-defined ER subdomains. Despite these limitations, our data underscore the purity of our ER membrane preparations, demonstrate a differential enrichment of ER subdomains (**Fig. 5**), and establish the lipid composition of the ER membrane (**Fig. 6**).

### The lipid composition of the ER from HEK293T cells

PC is by far the most abundant lipid class in the mammalian ER (**Fig. 6A**) in contrast to the ER from *S. cerevisiae*, where PC, PE and PI lipids contribute almost equally to the ER lipidome (Reinhard *et al*, 2024). The mammalian ER is composed of glycerophospholipids with short and mono-unsaturated fatty acyl chains, while the levels of cholesterol and sphingolipids are low (**Fig. 6**). Extrapolating from model membrane systems, this composition should result in loose lipid packing and render the ER membrane highly compressible. This may be functionally relevant as most membrane proteins are inserted into the membrane at the level of the ER by dedicated insertases that locally distort the lipid bilayer (Pleiner *et al*, 2020; Hegde & Keenan, 2021; Wu & Rapoport, 2021; McDowell *et al*, 2023). The ER membrane must accept, accommodate, and fold a large spectrum of membrane proteins differing in shape, hydrophobicity, and hydrophobic thickness (Hegde & Keenan, 2021; Renne & Ernst, 2023). Moving the ER membrane away from its biophysical optimum is expected to interfere with at least some of these crucial functions. In fact, membrane protein insertion into ER-derived microsomes is blocked by ramping up membrane cholesterol (Nilsson *et al*, 2001; Brambillasca *et al*, 2005), while increased lipid saturation and cholesterol in the ER causes lipotoxicity, associated with ER stress and activation of the unfolded protein response (Listenberger *et al*, 2003; Ariyama *et al*, 2010; Volmer *et al*, 2013; Halbleib *et al*, 2017; Piccolis *et al*, 2019; Radanović & Ernst, 2021). Hence, it is not surprising that the biophysical properties of the ER membrane and its sterol level are tightly controlled (Radhakrishnan *et al*, 2008; Brown *et al*, 2018; Harayama & Riezman, 2018; Covino *et al*, 2018; Radanović & Ernst, 2021)

Our lipidomic data align almost perfectly with a recent, systematic analysis of the organelle lipid compositions from HeLa cells using an alternative immunoisolation protocol (Foged *et al*, 2025). Like in HEK293T cells, PC is the most abundant lipid class in the ER, while the cholesterol and sphingolipid leves are low. The finding that ether PC (PC O-) and ether PE (PE O-) enrich in different compartments (Foged *et al*, 2025) is fully consistent with our observations on ether lipids in the ER: PC O- is enriched in the ER, while PE O- is depleted (**Fig. 6A, E**).

Nevertheless, there are also differences between the reported ER lipid compositions from HeLa and HEK293T cells. Particularly noteworthy are the even lower levels of SM (≈1 mol%) and cholesterol (7.8 mol%) in the ER of HeLa cells, contrasted by the higher levels of PC(>60 mol%) and Cer (≈0.43 mol%) (Foged *et al*, 2025) (**Fig. 6A-B, E-F**). These differences are not a mere consequence of different lipid class abundancies in the cell. HeLa cell lysate contains higher levels of PC (41 mol%), cholesterol (27 mol%), and SM (6.3 mol%) (Foged *et al*, 2025) than HEK293T lysates with only 37 mol% PC, 23 mol% cholesterol, and 2.7 mol% SM (**Suppl. Fig. S4A-B, E-F**). If HEK293T indeed establish more shallow lipid gradients along the secretory pathway than HeLa cells or if differences in cell cultivation, sample preparation (e.g. cell disruption, ER isolation) and lipid quantification affect the reported ER lipidomes remains to be established. Overall, both studies provide overwhelmingly consistent, deep insight into the lipid composition of the mammalian ER and pave way toward establishing more realistic ER membrane models for biochemical reconstitutions, biophysical characterization, molecular dynamics simulations.

### Transmembrane helices in the ER are more polar than in the plasma membrane

Using our newly developed tool, HelixHarbor, we extracted and analyzed the properties and composition of single-pass ER proteins TMHs for a comparison to TMHs in Golgi apparatus and the plasma membrane (**Fig. 4; Suppl. Fig. S3**). Expectedly, TMHs in the ER and the Golgi apparatus are shorter (**Suppl. Fig. S3A**) featuring larger sidechains (**Fig. 4B, D**) than plasma membrane TMHs, in which the accessible surface area is asymmetrically distributed (Sharpe *et al*, 2010; Lorent *et al*, 2017, 2025). Most striking, however, was the observation that the TMHs of ER proteins are significantly more polar than those in the plasma membrane (**Fig. 4C**). This increased polarity is matched by the surrounding lipids, considering that lipids with mono-unsaturated acyl chains, which are enriched in the ER, are more polar than lipids with saturated acyl chains (**Fig. 4C-D, G-H**) (Marsh, 2001; Zhanghao *et al*, 2020). Consistently, we observe that Cer lipids in the ER contain more hydroxylations than those in the whole cell lysate (**Suppl. Fig. S5A, B**). We can only speculate about the physiological relevance or purpose of an increased polarity of the ER membrane (**Fig. 4C**). It is likely that it contributes to membrane protein insertion and folding by lowering the activation barrier for moving polar protein portions through a distorted bilayer and by lowering the propensity of transmembrane domains with polar patches to collapse pre-maturely upon assembling macromolecular complexes.

Overall, we find a remarkable concordance of lipid and transmembrane helix features throughout our bioinformatic dataset and the Uniprot database (Lorent *et al*, 2025). Side chain bulkiness as measured by the accessible surface area (**Fig. 4D**) correlates inversely with lipid packing: The ER is composed of loosely packed lipids and hosts transmembrane domains with larger accessible surface areas. The plasma membrane, on the contrary, is highly asymmetric with loosely packed lipids in the cytosolic leaflet and tightly packed lipids in the extracellular membrane leaflet (Lorent *et al*, 2020), which is also reflected by the asymmetric distribution of bulky and small residues interfacing predominantly with the cytosolic and the extracellular leaflet of the plasma membrane, respectively (Sharpe *et al*, 2010; Lorent *et al*, 2020, 2025). Notably, there is robust experimental evidence that such protein-lipid concordance can contribute to the sorting of membrane proteins at various stations along the secretory pathway (Lorent *et al*, 2020; Castello-Serrano *et al*, 2024, 2025), hence underscoring co-evolution of transmembrane proteins and lipids based on their biophysical properties.

Zooming into the single amino acid level, we found fewer positively charged juxtamembrane residues in ER proteins compared to the proteins from the Golgi apparatus or the plasma membrane (**Fig 4.E-G**). It is tempting to speculate that this is another paradigm of protein-lipid concordance as the lower density of positively charged residues in ER membrane proteins is mirrored by the lower density of anionic lipids in the cytosolic leaflet of the ER in contrast to membranes of the late secretory pathway, where anionic lipids are actively enriched in the cytosolic leaflet (Bigay & Antonny, 2012; Holthuis & Menon, 2014). While C-terminal di-lysine motifs support a localization of transmembrane proteins to the ER (Jackson *et al*, 1990), a systematic characterization of the role of juxtamembrane residues in protein trafficking is still lacking. By automating the computation of physicochemical features such as surface area, bulkiness, and hydrophobicity, HelixHarbor provides a robust, reusable pipeline for future investigations into membrane protein properties.

In summary, mammalian MemPrep provides a comprehensive and quantitative view on the composition of the ER membrane. While our work does not exclude the possibility that local, specialized lipid environments may exist, we provide evidence that the lipid composition of ER tubules does not differ substantially from the rest of the ER. This work lays the foundation for generating more realistic ER membrane models.

## Materials and Methods

### Plasmid construction for retroviral transduction of HEK293T cells

Lists of oligonucleotides and primers used for genetic modification are provided (**Supplementary tables 2 and 3**). For the generation of pBABE-REEP5-3xFLAG-HRV3C-myc, the REEP5 gen was amplified by PCR from HEK293T cDNA using the primer pair pBABE-REEP5 fwd and REEP5-Tag rv. The tag was amplified by PCR from pRE866 (Reinhard *et al*, 2024) using the primers REEP5-Tag fwd and Tag-pBABE rv. Both PCR products were inserted into EcorRI linearized pBABE-puro by Gibson assembly (TakaraBio 638947). For the generation of pBABE-3xFLAG-HRV3C-myc-Sec61β, HRV3C-myc-Sec61β was synthesized (Invitrogen) and was inserted into p3XFLAG-CMV-24 (Sigma Aldrich). 3xFLAG-HRV3C-myc-Sec61β was amplified by PCR using the primers pBABE-Tag fwd and Sec61b-pBABE rv and was inserted into EcorRI linearized pBABE-puro by Gibson assembly. The full primary proteins sequence encoded by the constructs is provided in the **Supplementary table 4.**

### Antibodies

A complete list of antibodies used for this study is provided in the **Supplementary table 5**.

### Cell cultivation

Human embryonic kidney cells (HEK293T) were cultivated in DMEM substituted with 4.5 g/l glucose, 4 mM L-glutamine, 1 mM pyruvate, 10% (v/v) fetal bovine serum (FBS), at 37°C and 5% CO_2_. All cell lines were regularly tested negative for mycoplasma infections.

### Generation of stable cell lines

pBABE-REEP5-3xFLAG-HRV3C-myc or pBABE-3xFLAG-HRV3C-myc-Sec61β were transfected together with plasmids encoding for Gag-Pol (Addgene #8449) and VSV-G (Addgene #8454) into HEK293 helper cells using polyethylenimin. After 48 h, the supernatant was collected, supplemented with Polybrene (8 µg/mL) and used to infect the HEK293T target cells for 16 h followed by 48 h incubation. The polyclonal population of infected cells was selected in full DMEM medium with Puromycin (1 µg/mL) for 72 h to yield a stable cell line. The near-endogenous expression level of the bait construct was confirmed by SDS-PAGE and immunoblotting using antibodies directed against the FLAG epitope (M2, F1804, Sigma-Aldrich) and REEP5 (Abcam, ab186755).

### Mammalian MemPrep

For each MemPrep replicate followed by lipidomics as described in this section, cells were cultivated in 24*15 cm dishes to 90% confluency yielding approximately 4.8×10^8^ cells. For proteomics experiments, only halve of this material was used for each replicate. Cells were harvested, washed with 1x phosphate buffered saline (PBS), and pelleted by centrifugation at 600 x g. All subsequent steps were performed at 4°C using pre-chilled buffers. Cells were split in eight equal portions (each corresponding to 3*15 cm dishes), and each resuspended in 2 mL hypertonic lysis buffer (15% sucrose w/v, 10 mM 4-(2-hydroxyethyl)-1-piperazineethanesulfonic acid)(HEPES) pH 7.4, 300 mM NaCl, 1 mM EDTA freshly supplemented with protease inhibitor cocktail from Roche), and transferred into a precooled tight fitting Dounce Homogenizer (total volume 5 mL). Cells were lysed by 15 strokes on ice. The lysate in eight separate tubes was centrifuged (1,000 x g, 10 min, 4°C) to remove unbroken cells and debris. Each supernatant was transferred into a new tube and treated with Benzonase nuclease at a final concentration of 25 units/mL final concentration to degrade nucleic acids and lower sample viscosity. After another round of centrifugation (10,000 x g, 10 min, 4°C) to remove mitochondrial fragments, the resulting supernatants was subjected to ultracentrifugation (100,000 x g, 60 min, 4°C) using a TLA120.2 rotor (Beckmann) to pellet crude microsomes enriched for ER-derived membranes. The supernatants containing soluble, cytosolic proteins were discarded. Each of the eight pellets was resuspended in 1 mL lysis buffer snap-frozen in liquid nitrogen and stored at −80 °C till further processing.

As a preparation for the immunoisolation, eight times 1600 µL of magnetic bead slurry (DynaBeads, protein G, 2.8 µm diameter, Invitrogen) were equilibrated in lysis buffer and each decorated with 6.4 µg of a monoclonal mouse anti-Flag antibody (Sigma-Aldrich, F1804) during 16 h incubation on an overhead rotor at 20 rpm.

Before mixing with the affinity matrix, each of the 8 crude microsome preparations were thawed on ice for 20 min and sonicated (Bandelin Sonopuls HD 2070; Sonotrode MS72) with 50% power output using 10 pulses of each 0.7 s (duty cycle 0.7). The sonicated microsomes were cleared from metal particles by centrifugation (3000 x g, 3 min, 4°C).

The antibody-decorated affinity matrix was mixed in eight portions with each 1 mL sonicated microsomes and incubated for two hours under constant agitation in an overhead rotor at 3 rpm to facilitate capture of the bait protein. Specifically bound membrane vesicles were washed away by rinsing the affinity matrix twice with 800 μL wash buffer (15% w/v sucrose, 50 mM HEPES pH 7.4, 300 mM NaCl, 1 mM EDTA and 600 mM urea; pH 7.4) and then twice with 2000 µL of lysis buffer (without protease inhibitors). Highly enriched ER vesicles were eluted by cleaving the bait tag in lysis buffer (15% sucrose in 50 mM HEPES, 300 mM NaCl, 1 mM EDTA, 1 mM DTT, pH 7.4) containing no protease inhibitor but 0.2 mg/mL affinity purified HRV3C protease.

After a two-hour incubation of the sample on an overhead rotation at 3 rpm, eluate was separated from the magnetic DynaBeads and was then centrifuged (78,000 rpm, 120 min, 4°C) in a Beckman TLA120.2 rotor. The supernatants were carefully removed, and the eight pellets were resuspended using a total of 300 µL of 1X PBS for subsequent lipidomic analyses, or in 1X PBS supplemented with 1% SDS for subsequent proteomic analyses. The final isolate was snap frozen in liquid nitrogen and stored at −80°C.

### Preparation of whole cell lysates

Cells were cultivated in 10 cm cell culture dishes to 90% confluency in DMEM supplemented with 10% FBS. Cells were harvested, washed with 1X PBS, and resuspended in 300 µL of lysis buffer (0.5% Triton X100, 20mM HEPES pH 7.2, 150mM NaCl, PIC added freshly). The cells lysate was carefully agitated through pipetting, incubated on ice for 30 min, and cleared by centrifugation (16000 x g, 5 min, 4°C).

### SDS-PAGE and immunoblotting

Protein concentrations were determined using the microBCA protein assay (Thermo Fisher Scientific #23235) according to the manufacturer’s recommendations. For SDS-PAGE, the sample was mixed with 5x reducing membrane sample buffer (8 M urea, 0.1 M Tris-HCl pH 6.8, 5 mM EDTA, 3.2% (w/v) SDS, 0.05% (w/v) bromphenol blue, 4% (v/v) glycerol, 4% (v/v) β-mercaptoethanol), incubated for 10 min at 65 °C and subjected onto 4–15% mini-PROTEAN TGX precast protein gels (Bio-Rad). Proteins were separated in 1X Tris/Glycine/SDS buffer at 300 V for 20 min. After separation, proteins were transferred by semi-dry Western blotting onto nitrocellulose membranes (Amersham Protean Premium 0.45 µm). Proteins of interest were detected using specific primary antibodies and fluorescent secondary antibodies (**see Supplementary table 5**) on a fluorescence imager (LI-COR, Odyssey DLx). Signal intensities on immunoblots were quantified using ImageStudioLite.

### Immunofluorescence and microscopy

Cells were seeded on Poly-L-Lysine-coated cover slips and cultivated for 24 h in full DMEM medium at 37°C and 5% CO_2_. The cover slips were flash frozen in liquid nitrogen for 30 s and incubated in prechilled Methanol for 20 min on ice. The cells were washed with 3 times with PBS and were then transferred in blocking buffer (2% BSA in PBS). After 1 h incubation at RT, Immunolabelling was performed in PBS with 2% BSA using a 1:500 dilution of a mouse anti-FLAG antibody and a rabbit anti-Calnexin antibody for 1 h at room temperature. After ten wash steps, fluorescence staining was performed using a 1:500 dilution of a goat anti-rabbit antibody Alexa555 (Invitrogen, A21428) and a goat anti-mouse antibody (Invitrogen, A11029) Alexa488 supplemented with 5 µg/mL Hoechst33342 (Thermo Fisher scientific, H1399) in PBS with 2% BSA for 1 h at room temperature. Coverslips were washed ten times in PBS and another five times in water and were then mounted on specimen slides using Fluoromount-G mounting medium (Invitrogen, 00–4958-02). Images were taken on the Leica Dmi8 fluorescent microscope with a x63/1.40 NA objective and a Leica DFC3000 G CCD camera. z-Stacks covering the whole sample were recorded to ensure that the entire ER network was covered. Deconvolution was performed with Huygens essential software (17.04.0p8 64b) using the standard settings of the deconvolution wizard. Average-intensity projections of the central three planes were created in ImageJ (Schindelin *et al*, 2012). The contrast was adjusted linearly to correct for staining variations. For the quantification of co-localizations, maximum projections of three focal planes from a Z-stack were generated in ImageJ using micrographs that were not deconvoluted. Regions of interest were defined for individual cells, and the Manders coefficient was determined using the BIOP JACoP plugin (Bolte & Cordelières, 2006).

### Sample preparation for proteomics

Reduction of disulfide bridges of cysteines containing proteins was performed with dithiothreitol (56°C, 30 min, 10 mM in 50 mM HEPES, pH 8.5). Reduced cysteines were alkylated with 2-chloroacetamide (room temperature, in the dark, 30 min, 20 mM in 50 mM HEPES, pH 8.5). Samples were prepared using the SP3 protocol (Hughes *et al*, 2014, 2019) and trypsin (sequencing grade, Promega) was added in an enzyme to protein ratio 1:50 for overnight digestion at 37°C. Next day, peptide recovery in HEPES buffer by collecting supernatant on magnet and combining with second elution wash of beads with HEPES buffer. Peptides were labelled with TMT10plex (Werner *et al*, 2014) or TMT6plex Isobaric Label Reagent (ThermoFisher) according the manufacturer’s instructions. Samples were combined and for further sample clean up an OASIS® HLB µElution Plate (Waters) was used. For pull-down samples, up to 10 µg of peptides were labelled using TMTsixplex™ reagent as previously described (Dayon *et al*, 2008). Briefly, 0.8 mg of TMT reagent was dissolved in 45 µL of 100% acetonitrile. Subsequently, 4 µL of this solution was added to each peptide sample, followed by incubation at room temperature for 1 hour. The labelling reaction was quenched by adding 4 µL of a 5% aqueous hydroxylamine solution and incubating for an additional 15 min at room temperature. Labelled samples were then combined for multiplexing, desalted using an Oasis® HLB µElution Plate (Waters) according to the manufacturer’s instructions, and dried by vacuum centrifugation.

Offline high-pH reversed-phase fractionation (Yang *et al*, 2012) was carried out using an Agilent 1200 Infinity high-performance liquid chromatography (HPLC) system, equipped with a Gemini C18 analytical column (3 μm particle size, 110 Å pore size, dimensions 100 x 1.0 mm, Phenomenex) and a Gemini C18 SecurityGuard pre-column cartridge (4 x 2.0 mm, Phenomenex). The mobile phases consisted of 20 mM ammonium formate adjusted to pH 10.0 (Buffer A) and 100% acetonitrile (Buffer B). The peptides were separated at a flow rate of 0.1 mL/min using the following linear gradient: 100% Buffer A for 2 min, ramping to 35% Buffer B over 59 min, increasing rapidly to 85% Buffer B within 1 minute, and holding at 85% Buffer B for an additional 15 min. Subsequently, the column was returned to 100% Buffer A and re-equilibrated for 13 min. During the LC separation, 48 fractions were collected and combined for a total of 6 fractions (pull-down samples) or 12 fractions (full proteome samples) for MS analysis.

### Mass spectrometric (MS) analysis for proteomics

An UltiMate 3000 RSLCnano LC system (Thermo Fisher Scientific) equipped with a trapping cartridge (µ-Precolumn C18 PepMap™ 100, 300 µm i.d. × 5 mm, 5 µm particle size, 100 Å pore size; Thermo Fisher Scientific) and an analytical column (nanoEase™ M/Z HSS T3, 75 µm i.d. × 250 mm, 1.8 µm particle size, 100 Å pore size; Waters). Samples were trapped at a constant flow rate of 30 µL/min using 0.05% trifluoroacetic acid (TFA) in water for 6 min. After switching in-line with the analytical column, which was pre-equilibrated with solvent A (3% dimethyl sulfoxide (DMSO), 0.1% formic acid in water), the peptides were eluted at a constant flow rate of 0.3 µL/min using a gradient of increasing solvent B concentration (3% DMSO, 0.1% formic acid in acetonitrile). For pull-down samples, the gradient was as follows: 2% to 8% in 6 min (min), 8% to 28% in 72 min, 28% to 38% in 4 min, 38-80% in 0.1 min, maintained at 80% B for 2.9 min and a re-equilibrated to 2% B for 5 min. For full proteome samples, the gradient was as follows: 2% to 8% in 4 min (min), 8% to 28% in 104 min, 28% to 40% in 4 min, 40-80% in 0.1 min, maintained at 80% B for 3.9 min and re-equilibrated to 2% B for 4 min.

For pull-down samples labelled with TMT6, peptides were introduced into an Orbitrap Fusion™ Lumos™ Tribrid™ mass spectrometer (Thermo Fisher Scientific) via a Pico-Tip emitter (360 µm OD × 20 µm ID; 10 µm tip, CoAnn Technologies) using an applied spray voltage of 2.4 kV. The capillary temperature was maintained at 275°C. Full MS (MS1) scans were acquired in profile mode over an m/z range of 375–1,500, with a resolution of 60,000 at m/z 200 in the Orbitrap. The maximum injection time was set to 50 ms, the AGC target limit was set to ‘standard’. The instrument was operated in data-dependent acquisition (DDA) mode, with MS/MS scans acquired in the Orbitrap at a resolution of 15,000. The maximum injection time was set to 54 ms, with an AGC target of 200%. Fragmentation was performed using higher-energy collisional dissociation (HCD) with a normalized collision energy of 36%. MS2 spectra were acquired in profile mode. The quadrupole isolation window was set to 0.7 m/z, and dynamic exclusion was enabled with a duration of 60 s. Only precursor ions with charge states 2–7 were included for fragmentation. For full proteome samples labelled with TMT10, the same settings as described above were applied except for the following parameters. Full MS resolution was set to 120,000, MS2 resolution was set to a resolution of 30,000. The maximum injection time was set to 94 ms.

### Proteomics data analysis

Raw files were converted into the mzML format using MSConvert from ProteoWizard, using peak picking, 64-bit encoding and zlib compression, and filtering for the 1000 most intense peaks. Files were then searched using MSFragger in FragPipe (18.0-build13) (Kong *et al*, 2017) against the FASTA databaseUP000005640_HomoSapiens_ID9606_20594entries_26102022_dl1101202 3 containing common contaminants and reversed sequences. The following modifications were included into the search parameters: Carbamidomethylation (C, 57.0215), TMT (K, 229.162932) as fixed modifications; Oxidation (M, 15.9949), Acetylation (protein N-terminus, 42.0106), TMT (peptide N-terminus, 229.162932) as variable modifications. For the full scan (MS1) a mass error tolerance of 10 PPM and for MS/MS (MS2) spectra of 20 PPM was set. For protein digestion, ‘trypsin’ was used as protease with an allowance of maximum 2 missed cleavages requiring a minimum peptide length of 7 amino acids. The false discovery rate on peptide and protein level was set to 0.01. The standard settings of the FragPipe workflow ‘Default’ were used. For the proteomics data analysis, the raw output files of FragPipe (protein.tsv files files) were processed using the R programming environment (ISBN 3-900051-07-0). Initial data processing included filtering out contaminants and reverse proteins. Only proteins quantified with at least 2 razor peptides (with Razor.Peptides >= 2) were considered for further analysis. 2182 proteins passed the quality control filters. In order to correct for technical variability, batch effects were removed using the ‘removeBatchEffect’ function of the limma package (Ritchie *et al*, 2015) on the log2 transformed raw TMT reporter ion intensities (’channel’ columns). Subsequently, normalization was performed using the ‘normalizeVSN’ function of the limma package (VSN - variance stabilization normalization) (Huber *et al*, 2002). Missing values were imputed with the ‘knn’ method using the ‘impute’ function from the Msnbase package (Gatto & Lilley, 2011). This method estimates missing data points based on similarity to neighboring data points, ensuring that incomplete data did not distort the analysis. Differential expression analysis was performed using the moderated t-test provided by the limma package (Ritchie *et al*, 2015). The model accounted for replicate information by including it as a factor in the design matrix passed to the ‘lmFit’ function. Imputed values were assigned a weight of 0.01 in the model, while quantified values were given a weight of 1, ensuring that the statistical analysis reflected the uncertainty in imputed data. Proteins were annotated as hits if they had a false discovery rate (FDR) below 0.05 and an absolute fold change greater than 2. Proteins were considered candidates if they had an FDR below 0.2 and an absolute fold change greater than 1.5. Gene ontology (GO) enrichment analysis of hits was performed using the ‘compareCluster’ function of the ‘clusterProfiler’ package (Yu *et al*, 2012), which assesses over-representation of GO terms in the dataset relative to the background gene set. Enrichment was conducted for the following GO categories: Cellular Component (CC), Molecular Function (MF), and Biological Process (BP). The analysis was performed using ‘org.Hs.eg.db’ as the reference database. The odds ratio (’odds_ratio’) for each GO term was calculated by comparing the proportion of genes associated with that term in the dataset (’GeneRatio’) to the proportion in the background set (’BgRatio’). An odds ratio greater than 1 indicates that the GO term is enriched in the dataset compared to the expected background.

### TMH analysis using HelixHarbor

HelixHarbor is a custom, open-source tool developed for analyzing transmembrane helices (TMHs) and associated topological regions in transmembrane proteins. The tool is freely hosted and accessible at https://service2.bioinformatik.uni-saarland.de/HelixHarbor/, and a detailed Dcoker installation guide is provided in **Supplementary File 1**. TMH/topology definition and sequence extraction: For single-sequence analysis, HelixHarbor uses TMbed to assign per-residue classes (e.g., transmembrane helix/strand, signal peptide, or other) (Bernhofer & Rost, 2022). Contiguous runs of the same class are collapsed into segments and reported with 1-based start/end coordinates, segment type, and segment length. For analyses based on UniProt accessions, HelixHarbor retrieves protein sequences and annotated features from UniProt records (UniProt Consortium, 2025). Segment sequences are extracted using the annotated coordinate ranges (including transmembrane and topological domain features). Where available, experimentally derived topology constraints can be incorporated via TOPDB-derived annotations (Tusnády *et al*, 2008). Physicochemical features: For each TMH segment, HelixHarbor computes per-segment averages (mean per residue) of residue-level scales. Surface area is derived from the maximum allowed solvent accessibility reference values reported by Tien *et al*. (Tien *et al*, 2013) (reported in Å^2^ per residue) and averaged across residues in the segment. Bulkiness is calculated from the Zimmerman bulkiness index (Zimmerman *et al*, 1968) (unitless index), averaged across residues. Hydrophobicity is computed using the Kyte–Doolittle hydropathy scale (Kyte & Doolittle, 1982) (unitless index), averaged across residues. The tool also supports user-supplied custom residue scales (two-column amino-acid-to-value tables), summarized as the mean value per residue per segment. Comparisons, statistics, and visualization: Distributions of per-helix feature values are visualized as kernel density estimates (KDEs). KDE curves are probability densities whose integrals equal 1; each data point corresponds to the mean per-residue score across one TMhelix. Group differences can be assessed using standard two-sample tests (Kolmogorov–Smirnov, Welch’s t-test, and Mann–Whitney U). Scientific computing and plotting are performed in the SciPy ecosystem (Virtanen *et al*, 2020) with figures rendered using Matplotlib (Hunter, 2007).

### Establishment of transmembrane hydrophobicity profile

Transmembrane domain sequences from human single-pass transmembrane proteins of the ER, the Golgi apparatus, and the plasma membrane were retrieved from a previous publication (Lorent *et al*, 2025). Hydrophobicity of amino acids was estimated by the hydrophobicity scale of Kyte-Dolittle which was normalized to unity (Kyte and Doolittle 1982). To determine TMD hydrophobicity profiles, a distance of 0.15 nm in between amino acids along the α-helix was assumed. First, sequences were aligned to the middle of the membrane plane and for each sequence, hydrophobicity for each amino acid was calculated. The average hydrophobicity of each amino acid per position was then determined for each organelle-specific set of TMHs.

### Lipid extraction for mass spectrometry lipidomics

Mass spectrometry-based lipid analysis was performed by Lipotype GmbH (Dresden, Germany) as described (Surma *et al*, 2021). Lipids were extracted using a chloroform/methanol procedure (Ejsing *et al*, 2009). Samples were spiked with internal lipid standard mixture containing: cardiolipin 14:0/14:0/14:0/14:0 (CL), ceramide 18:1;2/17:0 (Cer), diacylglycerol 17:0/17:0 (DAG), hexosylceramide 18:1;2/12:0 (HexCer), lyso-phosphatidate 17:0 (LPA), lyso-phosphatidylcholine 12:0 (LPC), lyso-phosphatidylethanolamine 17:1 (LPE), lyso-phosphatidylglycerol 17:1 (LPG), lyso-phosphatidylinositol 17:1 (LPI), lyso-phosphatidylserine 17:1 (LPS), phosphatidate 17:0/17:0 (PA), phosphatidylcholine 15:0/18:1 D7 (PC), phosphatidylethanolamine 17:0/17:0 (PE), phosphatidylglycerol 17:0/17:0 (PG), phosphatidylinositol 16:0/16:0 (PI), phosphatidylserine 17:0/17:0 (PS), cholesterol ester 16:0 D7 (CE), sphingomyelin 18:1;2/12:0;0 (SM), triacylglycerol 17:0/17:0/17:0 (TAG) and cholesterol D6 (Chol). After extraction, the organic phase was transferred to an infusion plate and dried in a speed vacuum concentrator. The dry extract was re-suspended in 7.5 mM ammonium formiate in chloroform/methanol/propanol (1:2:4; V:V:V). All liquid handling steps were performed using Hamilton Robotics STARlet robotic platform with the Anti Droplet Control feature for organic solvents pipetting.

### Mass spectrometry (MS) data acquisition for lipidomics

Samples were analyzed by direct infusion on a QExactive mass spectrometer (Thermo Scientific) equipped with a TriVersa NanoMate ion source (Advion Biosciences). Samples were analyzed in both positive and negative ion modes with a resolution of Rm/z=200=280000 for MS and Rm/z=200=17500 for MSMS experiments, in a single acquisition. MSMS was triggered by an inclusion list encompassing corresponding MS mass ranges scanned in 1 Da increments (Surma et al. 2015). Both MS and MSMS data were combined to monitor CE, Chol, DAG and TAG ions as ammonium adducts; LPC, LPC O-, PC and PC O-, as formiate adducts; and CL, LPS, PA, PE, PE O-, PG, PI and PS as deprotonated anions. MS only was used to monitor LPA, LPE, LPE O-, LPG and LPI as deprotonated anions, and Cer, HexCer, and SM as formiate adducts and cholesterol as ammonium adduct of an acetylated derivative (Liebisch *et al*, 2006).

### Data analysis and post-processing for lipidomics

Data were analyzed with in-house developed lipid identification software based on LipidXplorer (Herzog *et al*, 2011, 2012). Data post-processing and normalization were performed using an in-house developed data management system. Only lipid identifications with a signal-to-noise ratio >5, and a signal intensity 5-fold higher than in corresponding blank samples were considered for further data analysis.

## Data Availability

The mass spectrometry proteomics data from this publication have been deposited to the ProteomeXchange Consortium via the PRIDE [http://www.ebi.ac.uk/pride] partner repository with the dataset identifier PXD075836 (Perez-Riverol et al, 2022). Other datasets are included in the source data file. The source code for HelixHarbor is available at https://github.com/mofty8/helixharbor-webserver.

## Autor Contributions

Aamna Jain: Resources; Data acquisition; Data curation; Formal analysis; Validation; Investigation; Visualization; Methodology; Writing-original draft; Writing-review and editing. Alexander von der Malsburg: Data acquisition; Validation; Resources; Methodology; Investigation; Visualization; Writing-review and editing. Claudia Götz: Resources; Validation; Investigation; Writing-review and editing. Mohamed Elmofty: Resources; Data curation; Formal analysis; Validation; Visualization; Methodology; Writing-review and editing. John Reinhard: Resources; Methodology; Writing-review and editing. Per Haberkant: Resources; Data curation; Formal analysis; Methodology; Writing-review and editing. Volkhard Helms: Resources; Data curation; Validation; Supervision; Funding Acquisition; Writing-review and editing. Joseph H. Lorent: Resources; Data curation; Formal analysis; Methodology; Writing-review and editing. Robert Ernst: Conceptualization; Data curation; Supervision; Funding acquisition; Validation; Visualization; Methodology; Writing—original draft; Project administration; Writing—review and editing.

## Supporting information

Supplementary Tabl 5

## Acknowledgements

This work was funded by the Deutsche Forschungsgemeinschaft (DFG, German Research Foundation) under Germanýs Excellence Strategy – EXC-3094 – 533751785 to RE and by the European Research Council under the European Union’s Horizon 2020 research and innovation program (grant agreement no. 866011) to RE. We acknowledge Julia Hach for excellent technical support throughout the project.

---

**Supplementary Figure S1:**
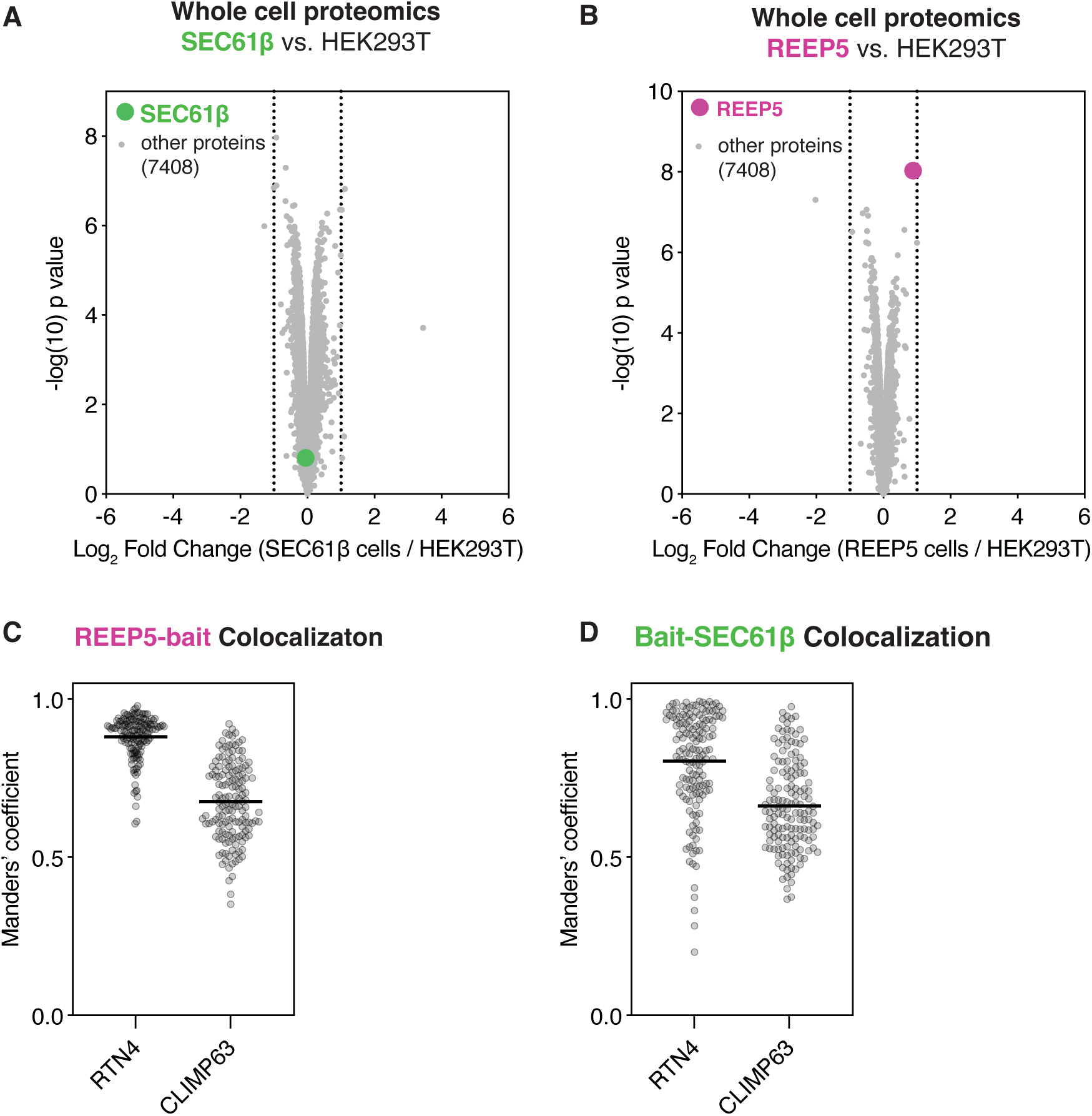
Impact of bait protein expression on HEK293T proteomes and correlation analysis of two MemPreps via the REEP5-bait and bait-Sec61β. **A:** Limma analysis of a non-targeted, TMT-labeling proteomics experiment performed with n=3 biological replicates comparing cell lysates from a bait-SEC61β expressing cell lines and that of HEK293T cells. **B:** Limma analysis of a non-targeted, TMT-labeling proteomics experiment performed with n=3 biological replicates comparing cell lysates from a REEP5-bait expressing cell lines and that of HEK293T cells. **C and D:** Quantification of the co-localization of the indicated bait proteins with the ER subdomain markers RTN4 (tubular ER) and CLIMP63 (ER sheets) (examples in Fig. 1E, F). Each data point represents the Manders’ coefficient of one individual cell. Per condition, 50 cells were quantified in three independent experiments (n = 150).

**Supplementary Figure S2:**
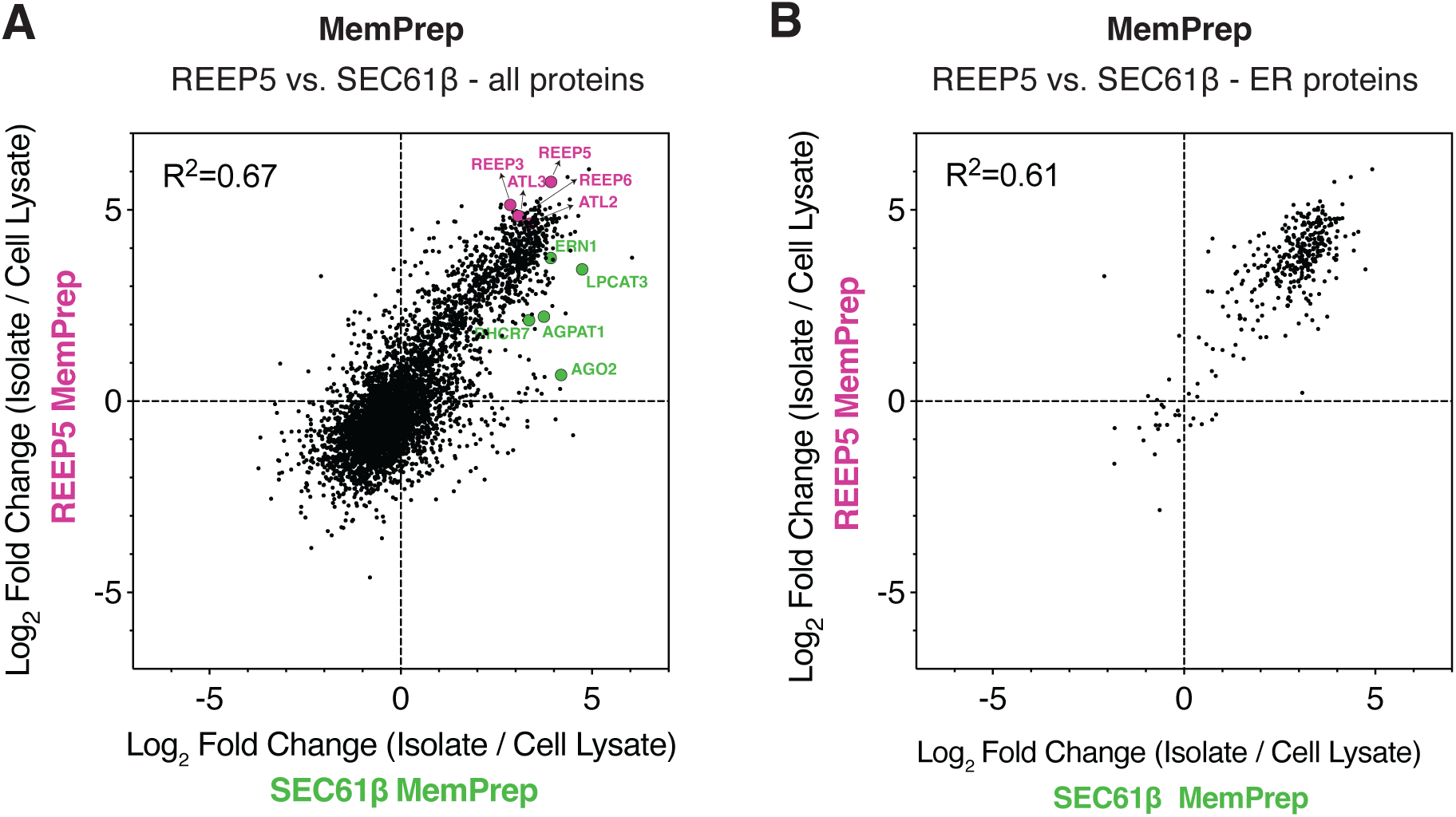
Correlation analysis of two MemPreps via the REEP5-bait and bait-Sec61β. **A:** The relative limma fold-change enrichment of proteins in the REEP5 and SEC61β MemPreps over the respective whole cell lysate were compared after non-targeted, TMT-labeling proteomics consistently detecting 4816 proteins. Plotting the limma log_2_-fold change in the isolate over the respective cell lysate from both MemPreps on two axes allows for a correlation analysis of the data. The correlation coefficient was determined by a linear fit forced through the origin. **B:** Correlation analysis as in (A) using a subset of the ER-annotated proteins in the MemPrep isolates. Each datapoint in (A) and (B) is from n = 3 biological replicates and from the same experiment as shown in Fig. 3.

**Supplementary Figure S3:**
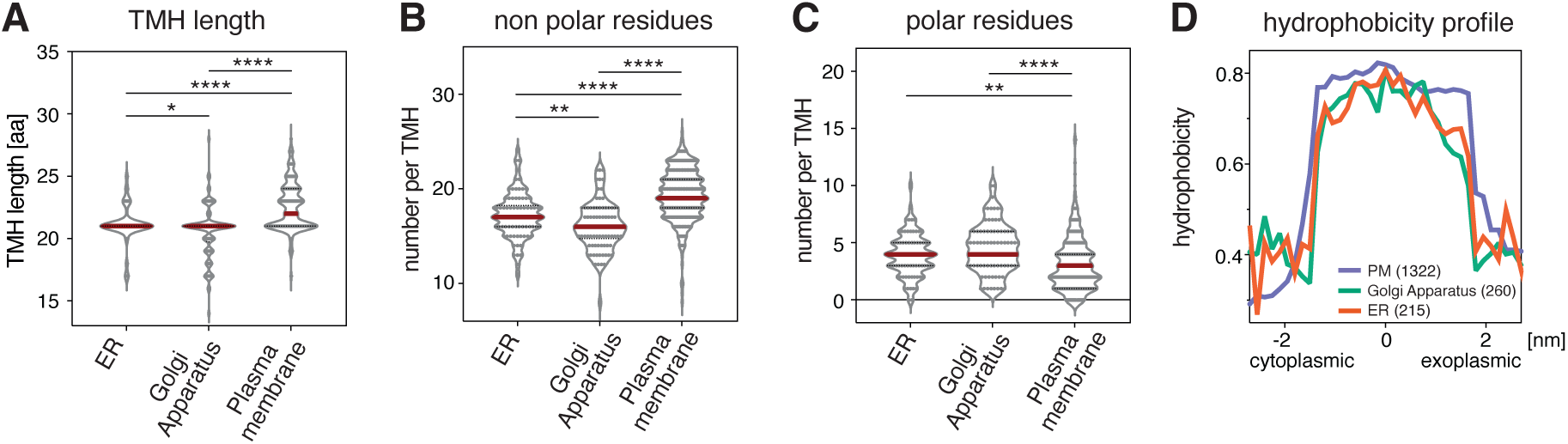
Systematic analysis of the transmembrane helix length and amino acid composition of single-pass ER proteins. **A:** The transmembrane domains of single-pass proteins from the ER (82), Golgi apparatus (90), and the plasma membrane (273) were systematically analyzed for their predicted TMH length. **B, C:** The number of non-polar (B) and polar (C) amino acids in each TMH of our dataset are plotted and sorted by the annotated, subcellular localization. The following amino acids (single letter code) were considered non-polar: L, I, V, A, M F, W, G, P; the following as polar: K, R, H, S, N, Q, T, C, Y, E, D. **D:** Average hydrophobicity profiles of all human single-pass TMDs from the ER (orange), the Golgi (green) and the PM (purple) along the membrane axis (cytoplasmic to exoplasmic) taken from (Lorent *et al*, 2025). Statistical test (A-C): unpaired T-tests with Welch’s correction *P≤0.05, **P≤0.01, ***P≤0.001, ****P≤0.0001.

**Supplementary Figure S4:**
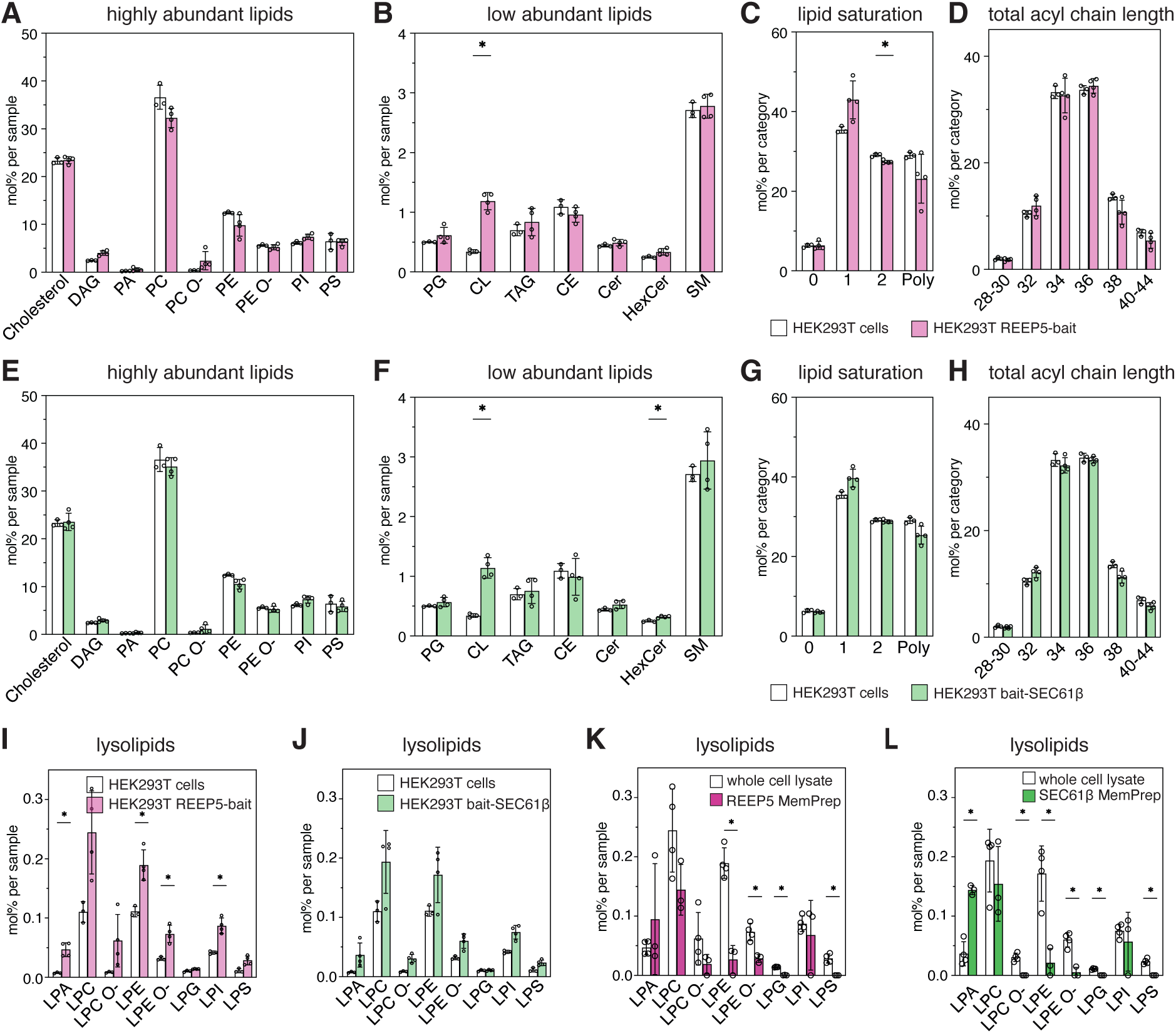
Expression of bait constructs has only a minor impact on the cellular lipidome. **A-H:** Quantitative lipidomics reveals the lipid class composition given as mol% of all identified lipids in the extract from a whole cell lysate from HEK293T WT cells, and cells expressing either the REEP5-bait or bait-SEC61β constructs. Except for cholesterol, each bar corresponds to a lipid class composed of several lipid species. **A, E:** *highly abundant lipids* - DAG: diacylglycerol, PA: phosphatidic acid, PC: phosphatidylcholine, PC O-: Phosphatidylcholine (-ether), PE: phosphatidylethanolamine, PC O-: Phosphatidylethanolamine (-ether), PI: phosphatidylinositol, PS phosphatidylserine). **B, F:** *low abundant lipids* - PG: phosphatidylglycerol, CL: cardiolipin, TAG: Triacylglycerol, CE: Cholesterol-ester, CE: Ceramide, HexCer: Hexosylceramide, SM. Sphingomyelin). **C, G:** Lipid unsaturation of whole cell lysates and the corresponding MemPrep isolate is given as the total number of double bonds in membrane glycerolipids except for CL (i.e. DAG, PA, PC, PC O-, PE, PE O-, PI, PS, PG) as mol% in this group. All lipids with more than two unsaturations in the lipid acyl chain are categorized as Poly-unsaturated lipids (Poly). **D, H:** The relative abundance of membrane glycerolipids with different total acyl chain lengths (calculated as the sum of both fatty acyl chains) in whole cell lysates and the corresponding MemPrep isolate are plotted in mol% of this category. **K, L:** Quantitative lipidomics reveals differences in lysolipid abundance in HEK293T whole cell lysates and the corresponding MemPrep isolates. The lysolipid composition is given as mol% of all identified lipids. Data information: Data from n = 4 biological replicates in (A-L) are represented as individual data points and as the mean ± SD. The data in (K-L) are from the same experiments as shown in Fig. 6. Statistical test: Multiple unpaired T-tests with Welch’s correction were performed by the Holm-Šídák method with a P-value threshold of 0.05. * represents qualitative significance.

**Supplementary Figure S5:**
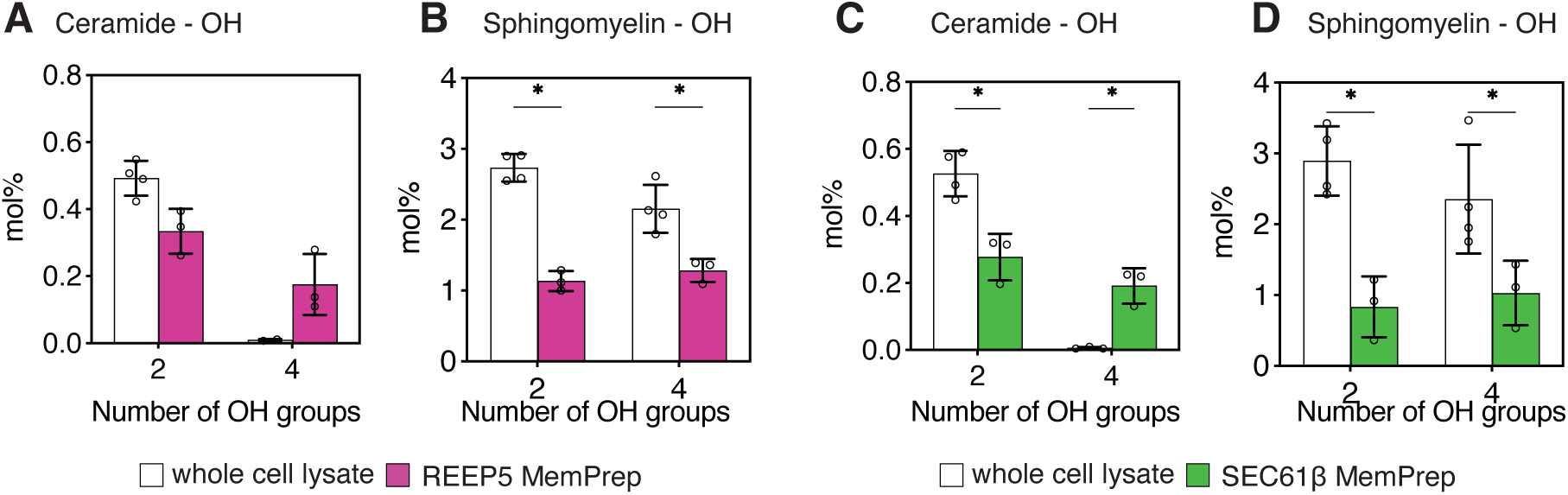
Ceramide hydroxylation, but not sphingolipid hydroxylation distinguishes the ER from the whole cell lipidome. **A-D:** Quantitative lipidomics of differentially hydroxylated sphingolipids in the whole cell lysate and MemPrep isolates. **A:** The level of differently hydroxylated Ceramide species is given in mol% of all identified lipids in the sample comparing a REEP5 MemPrep isolate with the corresponding whole cell lysate. **B:** The level of differently hydroxylated SM species is given in mol% of all identified lipids in the sample comparing a REEP5 MemPrep isolate with the corresponding whole cell lysate. **C:** The level of differently hydroxylated Ceramide species from SEC61β MemPreps and the corresponding cells are given in mol% and plotted as in (A) **D:** The level of differently hydroxylated SM species for SEC61β MemPreps and the corresponding cells are given in mol% and plotted as in (B). Data information: Data from n = 3 biological replicates in (A-D) are represented as individual data points and as the mean ± SD. Statistical test: Multiple unpaired T-tests with Welch’s correction were performed by the Holm-Šídák method with a P-value threshold of 0.05. * Represents qualitative significance.

**Supplementary table 1:** Total protein and total lipid yields of MemPrep procedures. The total protein and lipid amounts are given as the mean of n = 3 replicates with the standard deviation. Note that the amount of lipids in the whole cell lysates and in the MemPrep insolates were determined in separate experiments.

**Supplementary table 2:** All oligonucleotides used for cloning are listed in a 5’ to 3’ direction.

**Supplementary table 3:** Plasmids used in the study are listed together with the origin.

**Supplementary table 4:** Primary amino acid sequence of the bait tags used in the study.

**Supplementary table 5:** Antibodies used for immunoblotting and immunofluorescence are listed together with their origin.

