## Supplemental Table 1 for "Mammalian MemPrep establishes the lipid composition of ER membranes in HEK293T cells"

**Supplementary table 1: Total protein and total lipid yields**

| **REEP5 MemPrep** | **Protein** | **Protein yield** | **Lipid** | **Lipid yield** |
| --- | --- | --- | --- | --- |
| whole cell lysate | 12.3 ± 0.2 mg | 100% | 5.7 ± 2.3 µmol | 100% |
| P100.000 | 820 ± 97 µg | 6.7% | n.d. | n.d. |
| MemPrep isolate | 16.2 ± 7.9 µg | 0.13% | 68.8 ± 53.8 nmol | 1.2% |
| **SEC61β MemPrep** | **Protein** | **Protein yield** | **Lipid** | **Lipid yield** |
| whole cell lysate | 10.5 ± 1.5 mg | 100% | 4.9 ± 2.5 µmol | 100% |
| P100.000 | 977 ± 444 µg | 9.4% | n.d. | n.d. |
| MemPrep isolate | 19.6 ± 2.9 µg | 0.19% | 10.2 ± 6.7 nmol | 0.2% |

The total protein and lipid amounts are given as the mean of n = 3 replicates with the standard deviation. Note that the amount of lipids in the whole cell lysates and in the MemPrep insolates were determined in separate experiments.
