## Supplemental Table 2 for "Mammalian MemPrep establishes the lipid composition of ER membranes in HEK293T cells"

**Supplementary table 2: Oligonucleotides**

| **Oligonucleotide** | **Sequence (5'-3')** |
| --- | --- |
| REEP5-Tag fwd | AAGAGCACCGGGGGAGGCGGGGGTGGA |
| Tag-pBABE rv | GTCGACCACTGTGCTGGCGCTACTTGTCATCGTCATCCTTGTAGTCGATGTCATGATCTTTATAATCACCGTC |
| pBABE-REEP5 fwd | CCAGTGTGGTGGTACGTAGGATGTCTGCGGCCATGAGG |
| REEP5-Tag rv | CGCCTCCCCCGGTGCTCTTCTTTTCTTCACC |
| HindIII-preTAG fwd | AGAGAGAAGCTTGGGTCAGGTCTGGAAGTTCTGTTC |
| Sec61beta-EcoRI rv | AGAGAGGAATCCCTACGAACGAGT |
| XbaI-Sec61beta fwd | AGAGAGTCTAGAATGGACTACAAAGACC |
| Tag-KpnI rv | AGAGAGGGTACCCTACGAACGAGT |
| pBABE-Tag fwd | CCAGTGTGGTGGTACGTAGGATGGACTACAAAGACCATG |
| Sec61b-pBABE rv | GGTCGACCACTGTGCTGGCGCTACGAACGAGTGTACTTG |
