## Supplemental Table 3 for "Mammalian MemPrep establishes the lipid composition of ER membranes in HEK293T cells"

**Supplementary table 3: Plasmids**

| **Plasmid** | **Source** |
| --- | --- |
| pBABE | Martin van der Laan lab |
| pcDNA3.1 | Invitrogen |
| pRE866 | Ernst lab (Reinhard *et al*, 2024) |
| pMKRQ-_pre_TAP-tag-Sec61β | Synthesized by Invitrogen |
| p3XFLAG-CMV-24 | Sigma-Aldrich |
| p3XFLAG-HRV3C-Myc-Sec61beta-CMV-24 | This study |
| pBABE- REEP5-Tag | This study |
| pBAEBE-Tag-Sec61beta | This study |

Reinhard J, Starke L, Klose C, Haberkant P, Hammarén H, Stein F, Klein O, Berhorst C, Stumpf H, Sáenz JP, *et al* (2024) MemPrep, a new technology for isolating organellar membranes provides fingerprints of lipid bilayer stress. *EMBO J* 43: 1653–1685
