## Supplemental Table 4 for "Mammalian MemPrep establishes the lipid composition of ER membranes in HEK293T cells"

**Supplementary table 4: Primary sequence of bait constructs**

Highlighted sequence elements are the **3xFLAG tag, 3C protease recognition site** and the **myc tag**.

| **Name and size of bait construct** | **Sequence (single letter code)** |
| --- | --- |
| **REEP5-bait (241 aa)** | MSAAMRERFDRFLHEKNCMTDLLAKLEAKTGVNRSFIALGVIGLVALYLVFGYGASLLCNLIGFGYPAYISIKAIESPNKEDDTQWLTYWVVYGVFSIAEFFSDIFLSWFPFYYMLKCGFLLWCMAPSPSNGAELLYKRIIRPFFLKHESQMDSVVKDLKDKAKETADAITKEAKKATVNLLGEEKKSTGGGGGG**EQKLISEEDL**GSG**LEVLFQGP**GSG**DYKDHDGDYKDHDIDYKDDDDK** |
| **bait-Sec61β (152 aa)** | M**DYKDHDGDYKDHDIDYKDDDDK**LGSG**LEVLFQGP**GSG**EQKLISEEDL**GGGGGGASMPGPTPSGTNVGSSGRSPSKAVAARAAGSTVRQRKNASCGTRSAGRTTSAGTGGMWRFYTEDSPGLKVGPVPVLVMSLLFIASVFMLHIWGKYTRS |
