## Supplementary Tabl 5 for "Mammalian MemPrep establishes the lipid composition of ER membranes in HEK293T cells"

**Supplementary table 5: Antibodies used for immunoblotting (IB) and immunofluorescence (IF).**

| **Antibody** | **Dilution** | **Source** |
| --- | --- | --- |
| Mouse anti-GAPDH | IB: 1:1000 | Abcam (ab8245) |
| Rabbit anti-REEP5 | IB: 1:1000 | Abcam (ab186755) |
| Rabbit anti-SEC61B | IB: 1:1000 | Proteintech (15087-1-AP) |
| Rabbit anti-Rtn4 | IF 1:500 | Abcam (ab47085) |
| Rabbit anti-CLIMP-63 | IF 1:500 | Abcam (ab302539) |
| Rabbit anit-TOMM22 | IB: 1:1000 | Abcam (ab179826) |
| Rabbit anti-Histone H3 | IB: 1:1000 | Cell Signaling (9715S) |
| Rabbit anti-Calnexin | IB: 1:1000  IF: 1:500 | Proteintech (10427-2-AP) |
| Mouse anti-SERCA2 | IB: 1:1000 | Invitrogen (MA3-919) |
| Rabbit anti-Na/K-ATPase | IB: 1:1000 | Abcam (ab76020) |
| Rabbit anti-EEA1 | IB: 1:1000 | Abcam (ab109110) |
| Mouse anti-FLAG | IB: 1:1000  IF: 1:500 | Sigma-Aldrich (F1804) |
| Goat anti-Mouse IgG IRDye 800CW | IB: 1:2000 | LI-COR Bioscience (926-32210) |
| Goat anti-Rabbit IgG IRDye 800CW | IB: 1:2000 | LI-COR Bioscience (926-32211) |
| Goat anti-Mouse IgG Alexa Fluor™ 488 | IF: 1:500 | Thermo Fisher Scientific (A11029) |
| Goat anti-Rabbit IgG Alexa Fluor™ 555 | IF: 1:500 | Thermo Fisher Scientific (A21428) |
