## Supplementary File 1 for "Mammalian MemPrep establishes the lipid composition of ER membranes in HEK293T cells"

### HelixHarbor Docker Installation Guide

This guide provides instructions to download, run, and update the HelixHarbor Docker container.

#### Prerequisites

- Ensure Docker is installed on your system. If not, follow the official Docker installation guide.

#### Download and Run Docker Container

1. Pull the latest version of the Docker image:

   ```
   docker pull mmofty/helixharbor:versionx.x
   ```

   This command downloads the latest version of the image from Docker Hub.

2. Run the Docker container with the following command:

   ```
   docker run --restart=always -d -p 5005:5005 mmofty/helixharbor:versionx.x
   ```

   This command does the following:

   - `--restart=always`: Ensures the container restarts automatically if it crashes or if the Docker daemon restarts.
   - `-d`: Runs the container in detached mode.
   - `-p 5005:5005`: Maps port 5005 on your host to port 5005 in the container.
   - `mmofty/helixharbor:versionx.x`: Specifies the 'x.x' version of the Docker image to use.

3. Verify that the container is running:

   ```
   docker ps
   ```

   You should see an entry for the `mmofty/helixharbor:latest` container in the output.

#### Update to the Latest Version

1. Pull the latest version of the Docker image:

   ```
   docker pull mmofty/helixharbor:latest
   ```

2. List the running containers to find the container ID:

   ```
   docker ps
   ```

   Note the container ID for the currently running `mmofty/helixharbor` container.

3. Stop and remove the existing container:

```
docker stop <container_id> && docker rm <container_id>
```

Replace <container_id> with the actual container ID you obtained from the previous step.

4. Run the updated Docker container:

```
docker run --restart=always -d -p 5005:5005 mmofty/helixharbor:latest
```

5. Verify that the updated container is running:

```
docker ps
```

You should see an entry for the `mmofty/helixharbor:latest` container in the output.

#### Additional Tips

- **Check Logs**: If you encounter any issues, you can check the container logs with:

```
docker logs <container_id>
```

- **Stop Container**: To stop the running container:

```
docker stop <container_id>
```

- **Remove Container**: To remove a stopped container:

```
docker rm <container_id>
```

- **List All Containers**: To see all containers (running and stopped):

```
docker ps -a
```

- **Remove Unused Images**: To free up space by removing unused Docker images:

```
docker image prune -a
```
